# Genetically encoded intrabody sensors illuminate structural and functional diversity in GPCR-β-arrestin complexes

**DOI:** 10.1101/651463

**Authors:** Mithu Baidya, Punita Kumari, Hemlata Dwivedi, Eshan Ghosh, Badr Sokrat, Silvia Sposini, Shubhi Pandey, Tomek Stepniewski, Jana Selent, Aylin C. Hanyaloglu, Michel Bouvier, Arun K. Shukla

**Affiliations:** Department of Biological Sciences and Bioengineering, Indian Institute of Technology, Kanpur 208016, India; Institute for Research in Immunology and Cancer (IRIC), Université de Montréal, Montreal, Quebec, H3T 1J4, Canada; Department of Biochemistry and Molecular Medicine, Université de Montréal, Montreal, Quebec, H3T 1J4, Canada; Institute of Reproductive and Developmental Biology, Department of Surgery and Cancer, Hammersmith Campus, Imperial College London, Du Cane Road, London, W12 0NN, UK; Research Programme on Biomedical Informatics (GRIB), Department of Experimental and Health Sciences of Pompeu Fabra University (UPF)-Hospital del Mar Medical Research Institute (IMIM), 08003 Barcelona, Spain

**Keywords:** G protein-coupled receptors (GPCRs), β-arrestins, cellular signaling, synthetic antibody, intrabody, allosteric modulator, biased agonism, trafficking

## Abstract

Interaction of β-arrestins (βarrs) upon agonist-stimulation is a hallmark of G protein-coupled receptors (GPCRs) resulting in receptor desensitization, endocytosis and signaling. Although overall functional roles of βarrs are typically believed to be conserved across different receptors, emerging data now clearly unveils receptor-specific functional contribution of βarrs. The underlying mechanism however remains mostly speculative and represents a key missing link in our current understanding of GPCR signaling and regulatory paradigms. Here, we develop synthetic intrabody-based conformational sensors that help us visualize the assembly and trafficking of GPCR-βarr1 complexes in cellular context for a broad set of receptors with spatio-temporal resolution. Surprisingly, these conformational sensors reveal a previously unappreciated level of diversity in GPCR-βarr complexes that extends beyond the current framework of affinity-based classification and phosphorylation-code-based interaction patterns. More importantly, this conformational diversity arising from spatial signature of phosphorylation sites manifests directly in the form of distinct functional outcomes, including even opposite contribution of βarrs in signal-transduction for different receptors. Taken together, these findings uncover that despite an overall similar interaction and trafficking patterns; critical structural and functional differences exist in βarr complexes for different GPCRs that define and fine-tune receptor-specific downstream responses.

---

β-arrestins (βarrs) are multifunctional adaptor proteins, which play a central role in regulation and signaling of G protein-coupled receptors (GPCRs), the largest family of cell surface receptors in our body (*1, 2*). βarrs are evenly distributed in the cytoplasm under basal condition, and upon agonist-stimulation, they translocate to the plasma membrane to interact with activated and phosphorylated receptors (*3*). Binding of βarrs to GPCRs at the plasma membrane results in termination of G-protein coupling and desensitization of receptors through a steric hindrance based mechanism (*4*). Subsequently, βarrs drive receptor clustering in to clathrin-coated pits, via its ability to bind receptor, the β-subunit of the adaptor protein 2 (AP2) and heavy chain of clathrin, resulting in receptor internalization (*5*). βarrs either dissociate from the receptors and re-localize back in the cytoplasm, or they traffic into endosomal vesicles, in complex with the receptors, a feature associated with kinetics of post-endocytic sorting and βarr-mediated signaling and has led to classification of GPCR/βarr complexes as Class A (transient association, preference for βarr2) and Class B (sustained association, equal associations with βarr1/2) (*6*). βarrs also play a pivotal role in GPCR signaling by nucleating the components of various MAP kinase cascades such as p38, ERK1/2 and JNK3 (*7*).

As agonist-induced βarr recruitment to GPCRs is a highly conserved phenomenon, it can be used as a surrogate of GPCR activation, and also as a readout of βarr activation resulting in receptor endocytosis and signaling. Currently, a number of approaches are in use to monitor GPCR-βarr recruitment which include FRET/BRET based assays (*8*), enzyme complementation methods (*9*) and TANGO assay (*10*). Each of these methods necessitates a significant engineering and modification of the receptor, the βarr, or both. Here, we develop a set of intrabodies, which can specifically recognize receptor-bound βarr1, and report agonist-induced GPCR-βarr1 interaction and subsequent trafficking with spatio-temporal resolution in cellular context. Surprisingly, these intrabody sensors also reveal a previously unanticipated level of conformational diversity in GPCR-βarr complexes, which goes beyond the current classification of Class A vs. B, and phosphorylation-code-based βarr recruitment, and also functionally linked to distinct patterns of ERK1/2 MAP kinase activation.

Agonist-induced receptor activation and phosphorylation are two major driving forces for βarr recruitment and subsequent functional outcomes (*11*). A phosphorylated peptide corresponding to the carboxyl-terminus of the human vasopressin receptor V2R (referred to as V2Rpp) has been used extensively as a surrogate to induce active βarr conformation *in-vitro* (*12–14*). Previously, a set of synthetic antibody fragments (Fabs) have been generated against V2Rpp-bound βarr1 which selectively recognize active conformation of βarr1 as induced by V2Rpp (*15*). As the first step towards developing these Fabs as potential sensors of βarr1 activation and trafficking, we measured the ability of these Fabs to recognize βarr1 in complex with activated and phosphorylated β2V2R (*6*). Here, we used a chimeric version of β2AR harboring the carboxyl-terminus of the V2R, referred to as β2V2R which interacts with βarrs strongly. We observed that these Fabs were selectively able to interact with β2V2R-βarr1 complex that is formed upon agonist-stimulation of cells (Figure S1A-D).

A key step in developing these Fabs into cellular sensors of βarr1 activation and trafficking is to express them in a functional form in the cytoplasm as intrabodies. We therefore converted the selected Fabs into ScFvs (single chain variable fragments) by connecting the variable domains of their heavy and light chains through a previously optimized flexible linker (*16*), and then expressed them in mammalian cells as intrabodies either with a carboxyl-terminal HA tag or YFP fusion (Figure 1A-C and Figure S2A). We observed robust expression of two of these intrabodies namely Intrabody30 (Ib30) and intrabody4 (Ib4) in HEK-293 cells while others displayed relatively weaker expression (Figure 1B-C and Figure S2A). We also observed a significant level of nuclear localization of the intrabodies but the underlying reasons for this is currently not apparent to us. Interestingly, previous studies using intrabodies against β2AR such as nanobody 80 also reported a similar nuclear localization of fluorescently-tagged intrabodies (*2*).

**Figure 1.**
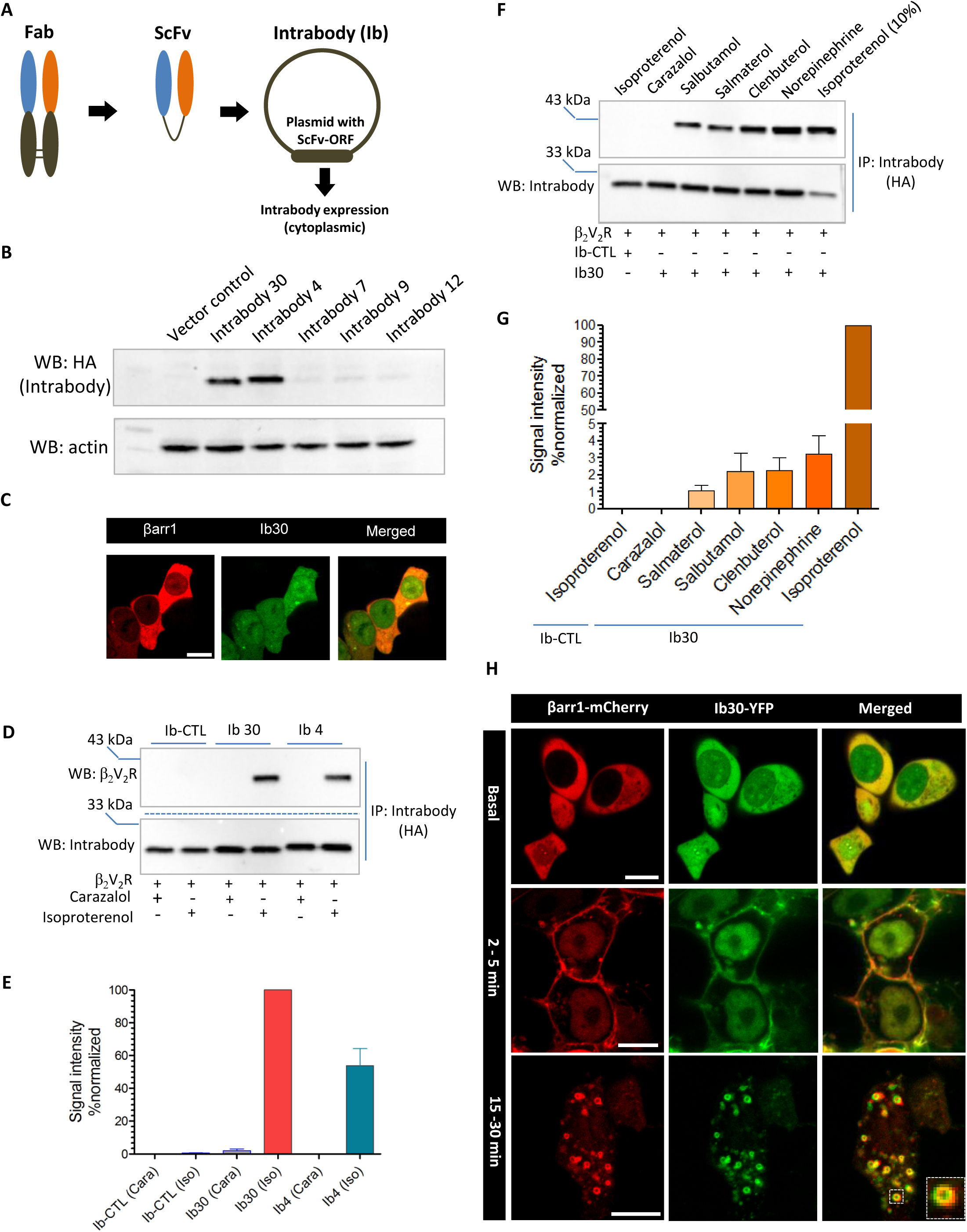
Generation and characterization of intrabody sensors for βarr1 recruitment and trafficking. **A**. Schematic representation of conversion of Fabs into ScFv format for intracellular expression (i.e. intrabodies). **B.** Expression analysis of intrabodies in HEK-293 cells as visualized by Western blotting. Lysate prepared from HEK-293 cells expressing the indicated intrabodies were separated on SDS-PAGE followed by visualization using anti-HA antibody. **C.** Cytoplasmic distribution of selected intrabodies as visualized by confocal microscopy. HEK-293 cells expressing βarr1-mCherry and YFP-tagged intrabodies were subjected to live cell imaging which revealed even distribution of intrabodies in the cytoplasm and nuclear localization. Scale bar is 10μm. **D**. Ability of intrabodies to recognize receptor-bound βarr1. HEK-293 cells expressing β2V2R, βarr1 and intrabodies were stimulated with either an inverse-agonist (carazolol) or agonist (Isoproterenol) followed by co-immunoprecipitation using anti-HA antibody agarose. The proteins were visualized by Western blotting using anti-Flag M2 antibody and anti-HA antibody. The bottom panel shows densitometry-based quantification of the data (average±SEM) from three independent experiments. **F**. The ability of Intrabody30 (Ib30) to recognize receptor-βarr1 complex mirrors the ligand efficacy. HEK-293 cells expressing β2V2R, βarr1 and intrabodies were stimulated with indicated ligands followed by co-IP and Western blotting as mentioned above. **G.** Densitometry-based quantification of the data (average±SEM) from three independent experiments. For isoproterenol, which yields maximal signal, only 10% of the total elution is loaded to avoid signal saturation. **H**. Intrabody30 reports agonist-induced βarr1 trafficking. HEK-293 cells expressing β2V2R, βarr1-mCherry and Ib30-YFP were stimulated with agonist (isoproterenol) for indicated time-points and the localization of βarr1 and Ib30 were visualized using confocal microscopy. Scale bar is 10 μm.

We then tested whether intrabodies maintain their ability to selectively recognize receptor-bound βarr1 in cellular context. For this, we first chose β2V2R because not only it forms a complex with βarr1 that is well-established to be recognized by Fab30 (*14, 16*), but also a set of ligands with broad efficacy profiles are available to allow in-depth characterization of intrabody sensors. We co-expressed β2V2R, βarr1 and selected intrabodies in HEK-293 cells, stimulated the cells with either an agonist or an inverse-agonist, and then immunoprecipitated intrabodies using carboxyl-terminal HA tag. We observed that both intrabodies i.e. Ib30 and Ib4 robustly recognized receptor-bound βarr1 exclusively upon agonist-stimulation (Figure 1D-E). Moreover, we also observed that the level of recognition of the receptor-βarr1 complexes by Ib30 mirrors the efficacy of the ligands (Figure 1F-G), which in turn further confirms the suitability of these intrabodies as a reliable sensor of βarr1 recruitment to the receptor and its activation.

We then co-expressed the β2V2R, βarr1-mCherry and YFP-tagged intrabodies in HEK-293 cells to monitor their trafficking patterns by confocal microscopy (Figure 1H and Figure S2B-D). As expected for β2V2R which behaves like a class B GPCR, we observed plasma membrane translocation of cytosolic βarr1 within a few minutes of agonist-stimulation, and interestingly, YFP-tagged intrabodies co-recruited with βarr1 to the plasma membrane and colocalized (Figure 1H, middle panel and Figure S2B). After prolonged agonist-exposure, βarr1 trafficked to the endosomal vesicles and once again, intrabodies exhibited co-localization with βarr1 (Figure 1H, lower panel and Figure S2B). Line-scan analysis further confirmed colocalization of intrabody sensors with βarr1 (Figure S2C-D). These observations taken together elucidate the ability of these intrabodies to faithfully report the trafficking pattern of βarr1 upon agonist-stimulation.

As these intrabodies were originally generated against V2Rpp-bound βarr1 conformation, we envisioned that they might be able to report βarr1 recruitment and trafficking for other chimeric GPCRs harboring the V2R-carboxyl-terminus. We therefore generated a set of chimeric GPCRs with V2R-carboxyl-terminus and measured the ability of Ib30 to recognize receptor-βarr1 complex by coimmunoprecipitation and trafficking of βarr1 by confocal microscopy (Figure 2A-X and supplementary Figures S3-S9). We observed that similar to β2V2R, Ib30 sensor robustly recognized receptor-βarr1 complexes for this broad set of chimeric receptors and also reported trafficking pattern of βarr1 with spatio-temporal resolution (Figure 2A-X and supplementary Figures S3-S9). These data establish that Ib30-YFP can be used as a generic tool to visualize agonist-induced βarr1 recruitment to the receptor and its subsequent trafficking in cellular context.

**Figure 2.**
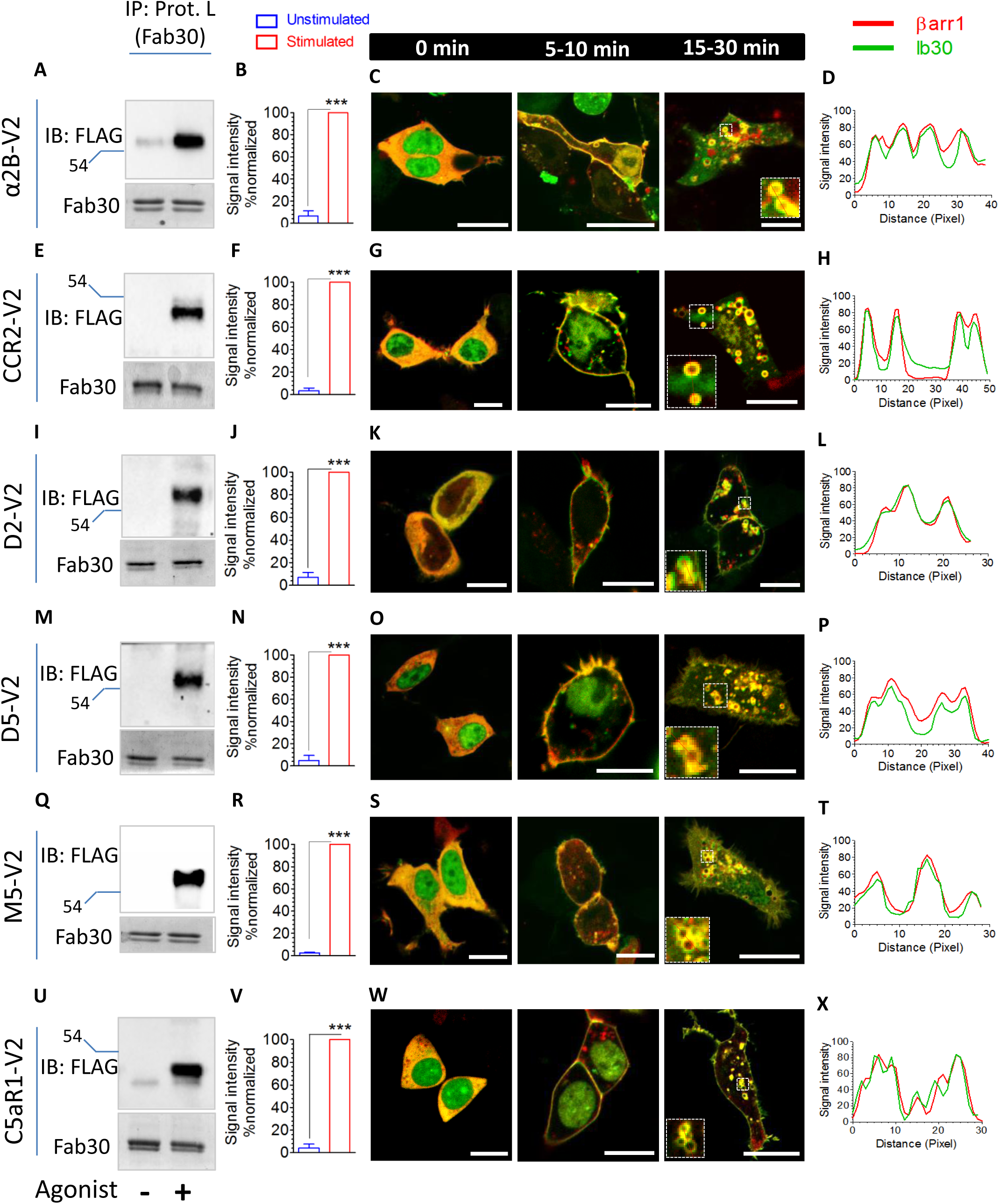
Intrabody30 as a generic sensor of βarr1 recruitment and trafficking for a set of chimeric GPCRs. HEK-293 cells expressing βarr1 (or βarr1-mCherry) and Ib30 (HA or GFP-tagged) were co-transfected with plasmids encoding N-terminally-FLAG tagged α2BV2R (**A**), CCR2V2R (**B**), D2V2R (**C**), D5V2R (**D**), M5V2R (**E**) and C5aR1V2R (**F**). Subsequently, cells were stimulated with respective agonists followed by co-immunoprecipitation using anti-HA antibody agarose. The proteins were visualized by Western blotting using anti-Flag M2 antibody and Simply Blue protein staining reagent (CBB) (Left panels). The right panels show the ability of Ib30 to report agonist-induced βarr1 trafficking for the corresponding GPCRs. HEK-293 cells expressing the respective receptor, βarr1-mCherry and Ib30-YFP were stimulated with agonist for indicated time-points and the localization of βarr1 and Ib30 were visualized using confocal microscopy (middle panels). Scale bar is 10μm. Right panels show line-scan analysis of images presented in the third sub-panel of confocal micrographs to demonstrate colocalization of βarr1-mCherry and IB30-YFP. Densitometry-based quantification of coIP data presented above (in panels B, F, J, N, R, V A – F) from three independent experiments normalized with respect to maximum signal for each receptor system (treated as 100%), and analyzed using One-way-ANOVA.

βarrs are capable of recognizing majority of GPCRs despite poorly conserved primary sequence of these receptors via agonist-induced receptor phosphorylation that drives βarr recruitment and activation. Therefore, we hypothesized that the intrabodies described here may recognize βarr1, and act as a sensor, for native GPCRs as well, considering that structural determinants for βarr1 recruitment and activation are likely to be conserved across the receptors. Accordingly, we next set out to test the intrabody30 on a broad set of native GPCRs using a combination of coimmunoprecipitation and confocal microscopy (Figure 3, Figure 4A-B, and Figure S10-12). We selected these receptors to encompass not only representatives of Class A and B (in terms of their interaction patterns with βarrs) but also a broad pattern of receptor phosphorylation codes proposed recently (*17*) (Figure 3A and Figure 4A). We started with V2R first and measured the ability of these intrabodies to recognize its complex with βarr1. We observed a pattern very similar to that of β2V2R in both, the the ability of Ib30 and Ib4 to form a complex with βarr1 via co-immunoprecipitation and co-traffic to the plasma and endomembrane compartments following receptor activation-(Figure 3B, 3D and Figure S10). V2R is classified as a class B GPCR based on its stable βarr recruitment pattern and co-internalization in complex with βarrs to endosomal vesicles. Expectedly, we observed a robust co-localization of V2R in endosomal vesicles with βarr1 and Ib30 which further confirms the ability of intrabodies to recognize receptor-βarr1 complexes and follow their endosomal translocation (Figure 3B and S13).

**Figure 3.**
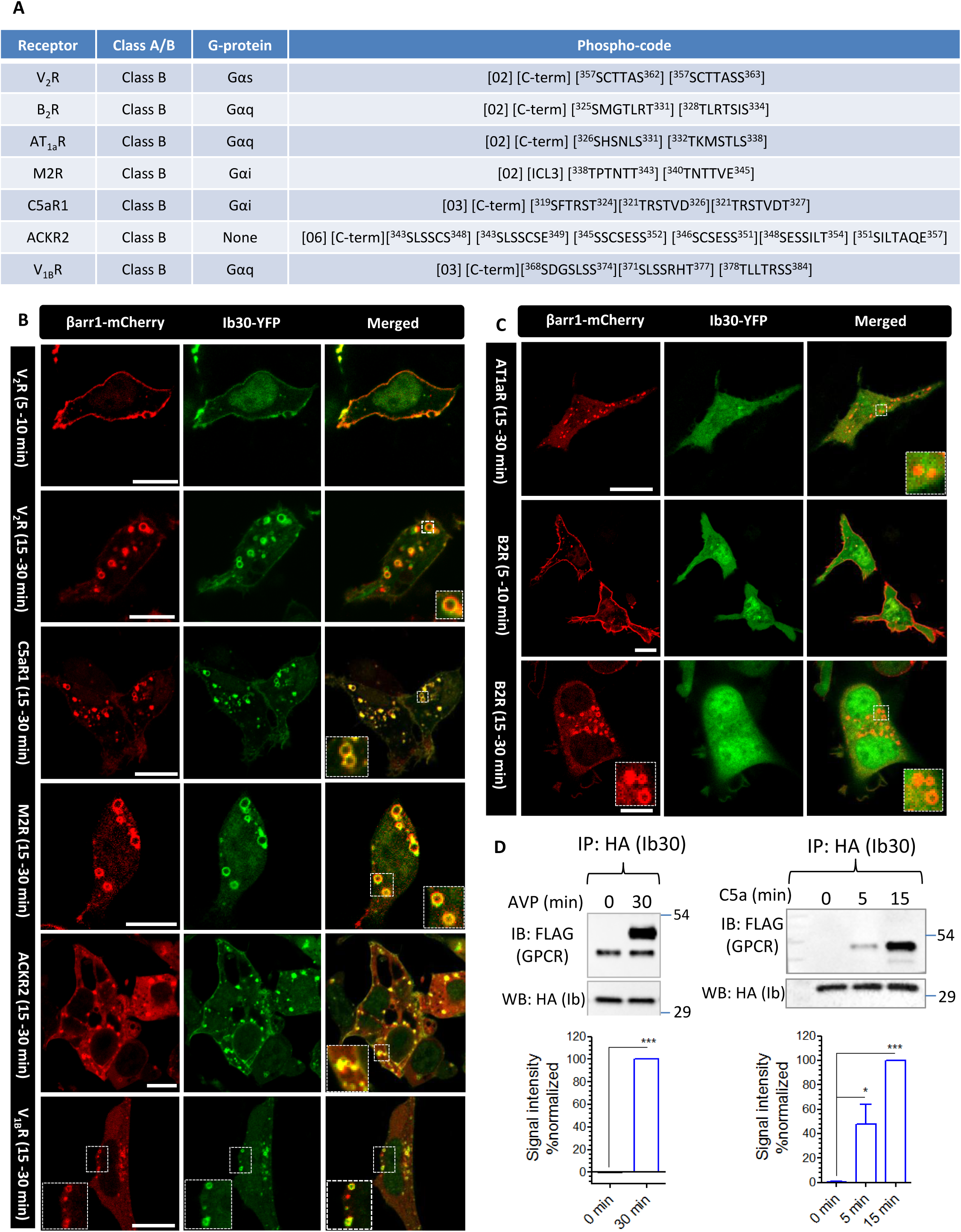
Intrabody30 sensor reveals conformational differences in βarr1 recruited to different class B GPCRs. **A**. G-protein-coupling preference and phospho-codes in a set of class B GPCRs (with respect to βarr recruitment pattern). Complete phospho-codes in these receptors are identified based on a recent study (*17*). **B**. Ib30-YFP sensor reports the recruitment and trafficking of βarr1 for several class B GPCRs such as V2R, C5aR1, ACKR2 and M2R, it does not recognize βarr1 upon activation of other class B GPCRs such as AT1aR and B2R as presented in panel **C.** For these experiments, HEK-293 cells expressing native GPCRs (as indicated in the respective panels), βarr1-mCherry and Ib30-YFP were stimulated with respective agonists for indicated time-points, and the localization of βarr1 and Ib30 were visualized using confocal microscopy. Scale bar is 10μm. **D**. The ability of Ib30 to recognize receptor-bound βarr1 is further confirmed by co-immunoprecipitation experiment. HEK-293 cells expressing either V2R or C5aR1, βarr1 and HA-tagged Ib30 were stimulated with agonist (100nM followed by co-immunoprecipitation using anti-HA antibody agarose. The proteins were visualized by Western blotting using anti-Flag M2 antibody and anti-HA antibody. The bottom panel shows densitometry-based quantification of the data (average±SEM) from three independent experiments normalized with respect to maximum reactivity (treated as 100%). Data are analyzed using One-Way-Anova with Bonferroni post-test (*p <0.05; ***p <0.001).

**Figure 4.**
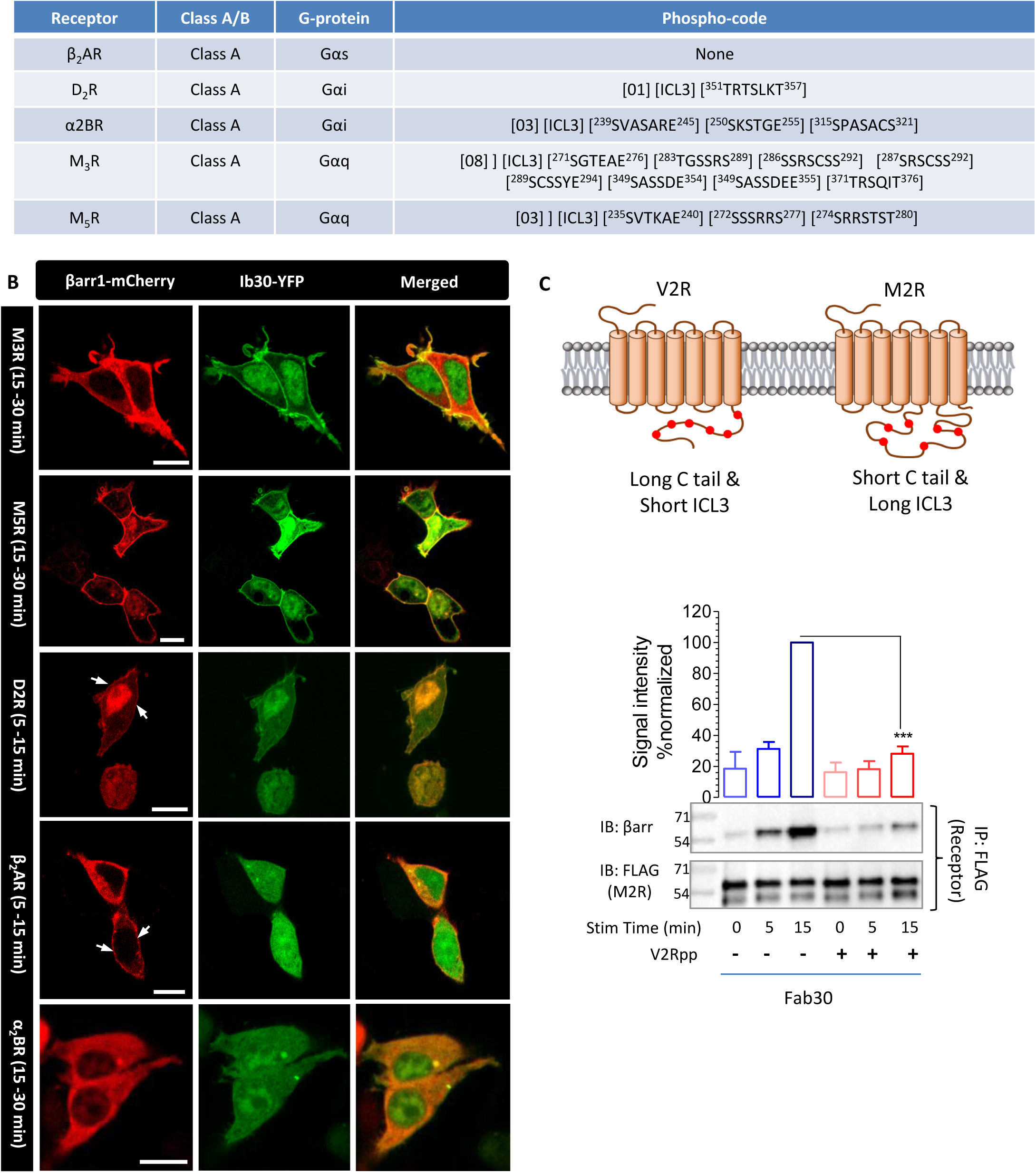
Intrabody30 sensor reveals conformational differences in βarr1 recruited to different class A GPCRs. **A.** G-protein-coupling preference and phospho-codes in a set of class A GPCRs (with respect to βarr recruitment pattern). Complete phospho-codes in these receptors are identified based on a recent study (*17*). **B**. Ib30-YFP sensor reports the recruitment and trafficking of βarr1 for M3R and M5R but it does not recognize βarr1 upon activation of other class A GPCRs such as β2AR and D2R. α2BR does not exhibit agonist-induced βarr1 recruitment despite having three potential phosphorylation codes in the 3^rd^ intracellular loops. For these experiments, HEK-293 cells expressing native GPCRs (as indicated in the respective panels), βarr1-mCherry and Ib30-YFP were stimulated with respective agonists for indicated time-points, and the localization of βarr1 and Ib30 were visualized using confocal microscopy. Scale bar is 10μm. **C**. The interaction of M2R with βarr1 is inhibited by V2Rpp suggesting an overlapping docking interface of its ICL3 with that of V2Rpp. The upper panel displays schematic representation of the V2R and M2R to underline the presence of phosphorylation sites (shown in red) in the carboxyl-terminus and ICL3, respectively. HEK-293 cells expressing M2R were stimulated with agonist followed by addition of purified βarr1 (with or without pre-incubation of V2Rpp) and Fab30. Subsequently, the receptor is co-immunoprecipitated using anti-Flag M1 agarose beads and proteins were visualized by Western blotting using anti-Flag M2 antibody and anti-βarr1 antibody. The bottom panel shows densitometry-based quantification of the data (average±SEM) from two independent experiments normalized with respect to maximum reactivity (treated as 100%). Data are analyzed using One-Way-Anova with Bonferroni post-test (**p <0.01).

Our results mentioned above for V2R strongly underscore the ability of intrabody sensors to report βarr1 recruitment and trafficking without any modification of the receptor or βarr1. As we had set-out to develop these intrabodies as specific sensors of βarr1 recruitment and trafficking, it is important that they do not exhibit any significant effect on heterotrimeric G-protein coupling. To confirm this, we measured agonist-induced cAMP response for V2R in presence of these intrabodies and observed that intrabodies did not significantly alter the kinetics or maximal cAMP response (Figure S14A-B). In addition, we also measured the effect of intrabodies on overall recruitment of βarr1 to the receptor, agonist-induced receptor endocytosis, and activation of ERK1/2 MAP kinases. As presented in Figure S15A-B, intrabodies slightly inhibited the recruitment of βarr1 to the plasma membrane but enhanced agonist-induced receptor endocytosis, as measured by BRET assays. They however did not significantly influence the kinetics of βarr1 localization in early endosomes (Figure 13B) or agonist-induced ERK1/2 phosphorylation (Figure S15C-F). Taken together, these data establish the suitability of Ib30 and Ib4 as a reliable sensor of receptor-βarr1 interaction, and subsequent trafficking of βarr1 in cellular context for native V2R.

We next tested the ability of Ib30 sensor to report the recruitment and trafficking of βarr1 for a broad set of native class B GPCRs including the complement C5a receptor (C5aR1), the atypical chemokine receptor 2 (ACKR2), the muscarinic M2 receptor (M2R), the vasopressin receptor sub-type 1b (V1bR), the angiotensin II type 1s receptor (AT1aR) and the bradykinin B2 receptor (B2R) (Figure 3B and Figure S11). These receptors not only couple to different sub-types of G-proteins but also harbor a diverse range of phosphorylation-codes in their carboxyl-terminus and the 3^rd^ intracellular loops (Figure 3A). We observed that Ib30 sensor worked efficiently for a number of these receptors including C5aR1, ACKR2, V1BR and M2R (Figure 3B and 3D). Interestingly however, it failed to recognize βarr1 upon stimulation of several others such as the AT1aR and B2R (Figure 3C) although these receptors exhibited significant level of βarr1 recruitment.

We also tested the ability of Ib30 sensor to report the recruitment and trafficking of βarr1 for a broad set of native class A GPCRs including the muscarinic M5 receptor (M5R), the muscarinic M3 receptor (M3R), the β2-adrenergic receptor (β2AR), the dopamine D2 receptor (D2R) and the α2-adrenergic receptor (α2BR) (Figure 4A-B and Figure S12). These receptors also couple to different sub-types of G-proteins and harbor a diverse range of phosphorylation-codes in their carboxyl-terminus and the 3^rd^ intracellular loops (Figure 4A). Similar to class B GPCRs mentioned above, we observed that Ib30 sensor robustly recognizes βarr1 for the muscarinic M5 receptor (M5R) and the muscarinic M3 receptor (M3R) but not for others. Taken together, the Ib30 reactivity pattern suggests that the conformation of βarr1 in complex with various receptors can be significantly different from each other despite comparable patterns of overall gross βarr1 recruitment. We note here that M3R and M5R did not induce any significant endosomal localization of βarr1 even after prolonged agonist-stimulation as expected for class A GPCRs.

An interesting observation here is the ability of Ib30 sensor to robustly recognize βarr1 upon stimulation of muscarinic receptors namely M2R, M3R and M5R. This is striking because these receptors, unlike V2R, possess relatively large 3^rd^ intracellular loop and very short carboxyl-terminus (Figure 4C). More importantly, they harbor the full phospho-codes exclusively in their 3^rd^ intracellular loops. The reactivity pattern of Ib30 suggests that these receptors, despite having phosphates in distinct receptor domains, are able to induce a conformation in βarr1 that allows recognition by Ib30. As mentioned earlier, the crystal structure of V2Rpp-βarr1 complex reveals the docking of V2Rpp on the N-domain of βarr1. Thus, it is tempting to speculate that these receptors may also be able to engage an identical, or at least similar, interface on βarr1 through their 3^rd^ intracellular loops harboring the phosphorylation sites. In order to test this hypothesis and further corroborate our findings, we measured the interaction of M2R with βarr1 in presence and absence of V2Rpp. As presented in Figure 4C, pre-incubation of V2Rpp with βarr1 robustly inhibits the interaction between M2R and βarr1, and therefore, suggest that M2R engages βarr1 through the N-domain interface similar to that engaged by V2Rpp. This is particularly intriguing as the first step in GPCR-βarr interaction is typically considered to involve the phosphorylated carboxyl-terminus followed by the core interaction involving the intracellular loops of the receptors. Considering the short carboxyl-terminus of muscarinic receptors typically devoid of any phosphorylation codes, coupled with much larger, phosphorylation code containing 3^rd^ intracellular loop (ICL3), it is tempting to speculate a distinct mode of receptor-βarr interaction compared to other GPCRs. In fact, a recent study on M1R using optical reporters suggests two distinct conformations of receptor-bound βarrs, which are also distinct from biphasic interaction observed for other prototypical GPCRs (*18*). Furthermore, it also raises the possibility that other receptors harboring short C-terminus but long ICL3 such as the dopamine D2 receptor and α-adrenergic receptors, may also display distinct binding conformations compared to other prototypical GPCRs.

The findings described up to this point have two important implications. First, they establish the utility of the intrabody sensors described here as a robust tool to visualize agonist-induced βarr1 recruitment and trafficking for a number of GPCRs. Second, they imply that different receptors induce distinct conformations in βarr1 even if they exhibit a similar pattern of gross-recruitment (e.g. V2R vs. B2R) and harbor a similar phospho-code signature in their carboxyl-terminus and 3^rd^ intracellular loops (Figure S16). This prompted us to ask if these distinct conformations of βarr1 for different receptors might have different functional consequences. In order to probe this possibility, we measured agonist-induced ERK1/2 MAP kinase activation for selected receptors and the role of βarr1 in this process (Figure 5A-B and Figure S17-18). As discussed earlier, one of the key contributions of βarrs in GPCR signaling is their ability to mediate agonist-induced phosphorylation and activation of ERK1/2 MAP kinases (*7*). We observed that knockdown of βarr1 yielded a significant reduction in ERK1/2 phosphorylation for V2R while it augmented the levels of ERK1/2 phosphorylation activated by GPCRs that did not recruit the arrestin intrabodies such as B2R and AT1aR (Figure 5A-B and Figure S17-18). These striking differences in the functional contribution of βarr1 in agonist-induced ERK1/2 activation suggests a potential link between the receptor-bound βarr1 conformation and different functional outcomes, and also provide a potential structural mechanism for how receptor-specific βarr conformations encode functional differences. An important implication of this data is that βarr1 conformation upon stimulation of B2R is competent to drive endocytosis but suppressive towards ERK1/2 phosphorylation.

**Figure 5.**
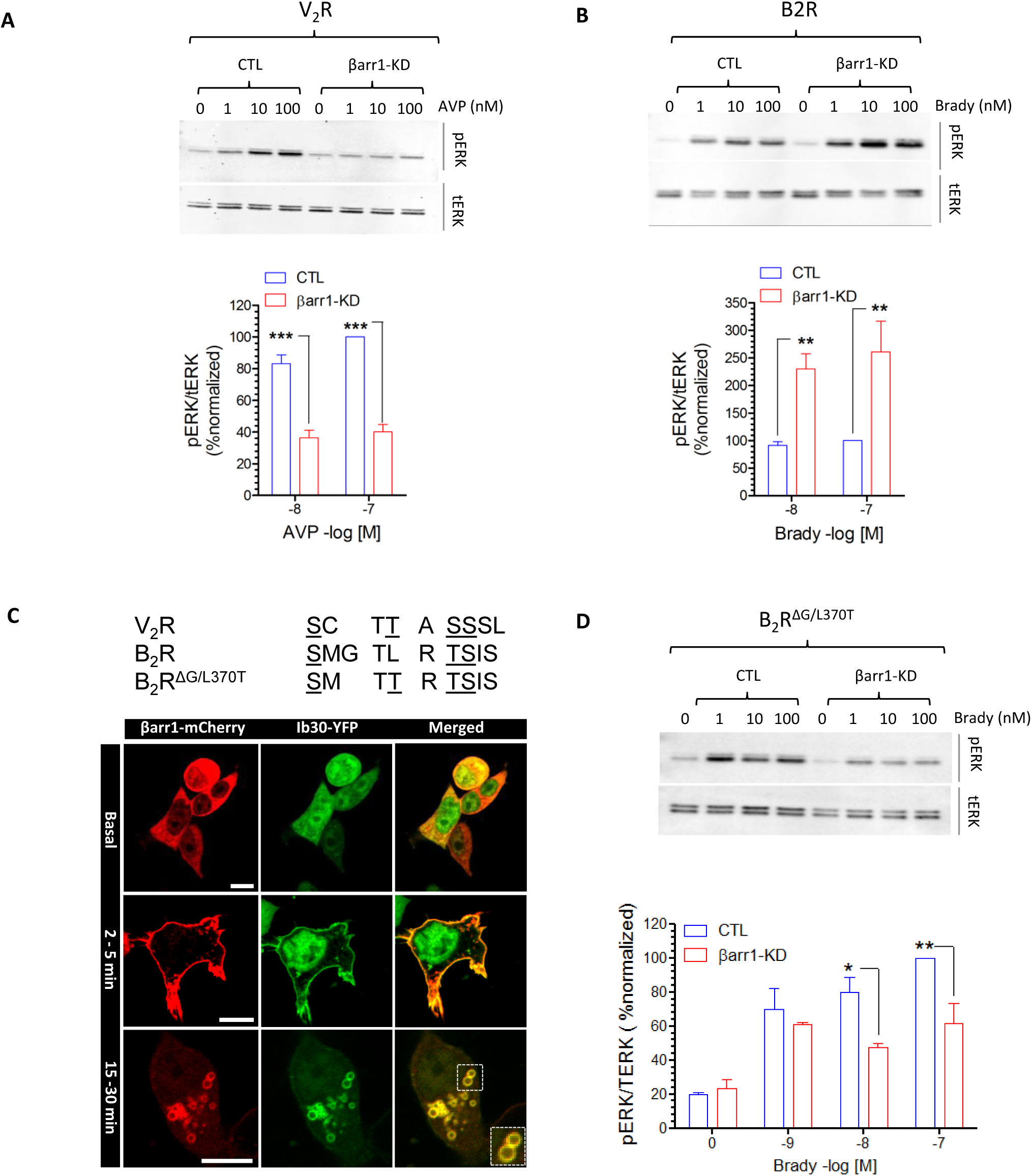
Distinct conformations of receptor-bound βarr1 are linked with different functional outcomes. Different conformations of receptor-bound βarr1 drive their distinct functional contribution in agonist-induced ERK1/2 MAP kinase phosphorylation for V2R and B2R. Agonist-induced phosphorylation of ERK1/2 in HEK-293 cells expressing either V2R (**A**) or B2R (**B**) in presence and absence of βarr1 knock-down are measured using Western blotting. Densitometry-based quantification of data from three independent experiments is presented as bar-graphs in the right panels normalized with respect to maximal signal under control condition (treated as 100%) and analyzed using One-way-ANOVA with Bonferroni post-test (**p <0.01). C. Comparison of spatial distribution of phosphorylation sites in V2R and B2R. The double mutant of B2R, referred to as B2R^ΔG/L370T^, is generated to mimic the spatial distribution of Ser/Thr in V2R. Confocal microscopy reveals robust recognition of βarr1 upon agonist-stimulation of B2R^ΔG/L370T^. HEK-293 cells expressing B2R^ΔG/L370T^, βarr1-mCherry and Ib30-YFP were stimulated with agonist (bradykinin, 1μM) for indicated time-points and the localization of βarr1 and Ib30 were visualized using confocal microscopy. Scale bar is 10μm. **D**. Knock-down of βarr1 robustly inhibits agonist-induced ERK1/2 phosphorylation for B2R^ΔG/L370T^, similar to V2R and in stark contrast with wild-type B2R. HEK-293 cells expressing B2R^ΔG/L370T^ in presence and absence of βarr1 knock-down were stimulated with indicated doses of bradykinin for 10 min followed by detection of phosphorylated ERK1/2 using Western blotting. Densitometry-based quantification of data from three independent experiments is presented as bar-graphs in the lower panel normalized with respect to maximal signal under control condition (treated as 100%) and analyzed using One-way-ANOVA with Bonferroni post-test (*p <0.05; **p <0.01).

As mentioned above, both V2R and B2R belong to class B GPCRs, in terms of βarr recruitment patterns, and they each harbor two full phosphorylation codes in their carboxyl-terminus. This suggests that distinct patterns of Ib30 reactivity and ERK1/2 phosphorylation arise elsewhere. A close comparison of the carboxyl-terminus of these two receptors revealed a difference in the spatial distribution of the phosphorylation sites (Figure 5C). This prompted us to generate a set of B2R mutants to resemble the pattern of phosphorylation in V2R (Figure S19A). These mutants exhibited comparable surface expression levels similar to wild-type B2R (Figure S19B). Out of these mutants, we did not observe any detectable reactivity of Ib30 for B2R^ΔG368^ and B2R^I374S^ although they robustly recruited βarr1 (Figure S20A-B). Interestingly however, we found robust reactivity of Ib30 in case of B2R^ΔG368+L374T^ and B2R^L374T^ (Figure 5C, Figure S19B). These findings reveal that spatial signature of receptor phosphorylation sites play a detrimental role in ensuing βarr1 conformation recognizable by Ib30. Even more strikingly, we discover that βarr1 plays a positive role in phosphorylation of ERK1/2 for the B2R^ΔG368+L374T^ and B2R^L374^ as the lack of βarr1 results in a substantial reduction (Figure 5D and Figure S19C-D). The pattern of ERK1/2 phosphorylation for B2R^ΔG368+L374T^ and B2R^L374^ is similar to that of V2R, and it is in stark contrast with the wild-type B2R, thereby linking the βarr1 conformation, as read by Ib30, with subsequent functional outcome. These striking observations indicate that βarr1 adopts a conformation upon stimulation of B2R^ΔG368+L374T^ and B2R^L374^ that is capable of supporting both, endocytosis and ERK1/2 phosphorylation, unlike the wild-type B2R.

In order to gain structural insights into the recognition of B2R mutants by intrabody30 sensor, we employed molecular dynamics simulation approach using the previously determined crystal structure of βarr1 in presence of V_2_Rpp. We carried out MD simulations with V2Rpp, B2Rpp and the mutant version of B2Rpp (derived from B2R^L374T^) (Figure S21A). We observed that in the B2Rpp mutant as well as V2Rpp, L360pT together with pT359 interacts with K294 in the lariat loop via strong electrostatic interactions (Figure S21B, blue bar plots). Such bifurcated contacts seem to stabilize the lariat loop preferential in conformational states belonging to cluster 1 (Figure S21C, blue lines in structure and blue bar plots). Cluster 1 overall resembles the conformation of the crystallized lariat loop in complex with Fab30 (pdb 4JQI) with an average rmsd of 1.9 Å. Such bifurcated linkage of the lariat loop to the phosphorylated receptor C-tail is lost in the B2Rpp WT as the non-polar L360 cannot establish an interaction with K294 (Figure S21B, red bar plot). As a consequence, the loop conformation is shifted downwards favoring cluster 2 (Figure S21C), red lines in structure and red bar plot). This yields a dramatic increase of average rmsd to 4.6 Å with respect to the crystallized lariat loop. It is likely that such conformational change of the lariat loop in the Fab30 binding interface contributes to a decreased Fab30 binding. Overall, MD data together with site directed mutagenesis suggest that the lariat loop in βarr1 may provide an important site for driving conformational differences in different GPCR-βarr1 complexes, at least as measured by intrabody30 reactivity.

Taken together, we develop intrabody-based sensors, which allow direct visualization of agonist-induced recruitment of βarr1 to a broad set of GPCRs and report βarr1 trafficking with spatio-temporal resolution. Interestingly, these Intrabody sensors reveal significant conformational diversity in GPCR-βarr1 complexes, which manifest in distinct contribution of βarr1 in agonist-induced ERK1/2 activation. Although interaction of βarrs is highly conserved across GPCRs, there are many instances of distinct functional contribution of βarrs in trafficking and signaling of different GPCRs despite having an overall similar recruitment and trafficking pattern (*19*). These examples suggest that all GPCR-βarr complexes formed in cells may not have identical functional abilities and recent studies have in fact started to provide evidence towards this notion (*20, 21*). Intrabody sensors developed here now offer direct evidence for distinct conformational signatures in βarr1, and therefore, offer important insights into the process of GPCR activation, trafficking and signaling. Our data with B2R mutants also underscore that βarr conformations supporting receptor endocytosis vs. ERK1/2 phosphorylation are likely to be distinct from each other. This has direct implications for the conceptual framework of biased-agonism at GPCRs aimed at designing novel therapeutic strategies.

## Acknowledgment

We thank the members of our laboratories for critical reading of the manuscript. The research program in our laboratory is supported by the DBT Wellcome Trust India Alliance (Intermediate Fellowship to A.K.S.—IA/I/14/1/501285), Department of Biotechnology, Government of India (Innovative Young Biotechnologist Award to A.K.S.—BT/08/IYBA/2014-3), LADY TATA Memorial Trust Young Researcher Award to A.K.S., Science and Engineering Research Board (SERB) (SB/SO/BB-121/2013), Council of Scientific and Industrial Research (CSIR) (37[1637]14/EMR-II). A.K.S. is an EMBO Young Investigator. AH is supported by the Genesis Research Trust (P73441) and the Biotechnology and Biological Sciences Research Council (BB/N016947/1, BB/S001565/1).

## Author’s contribution

MB and PK carried out most of the experiments related to intrabody generation, characterization, confocal microscopy, co-immunoprecipitation and ERK1/2 phosphorylation; EG performed the ERK1/2 phosphorylation for V2R and contributed in generation of receptor mutants; SP carried out the ERK1/2 phosphorylation experiment together with PK; BS performed the BRET experiments under the supervision of MB; SS carried out the 3-colour confocal and endosomal co-localization experiments under the supervision of AH; SS measured the colocalization of V2R,βarr1 and Ib30 under the supervision of AH; BS carried out the BRET experiments under the supervision of MB; AKS supervised the overall project execution and management.

## Materials and methods

### Cell Culture and Transfection

HEK293 cells (ATCC) were maintained in DMEM containing 10% FBS and penicillin/streptomycin (100 U/mL) at 37°C in 5%CO2. Transient transfections of DNA were performed with Lipofectamine 2000 (Life Technologies) and cells were assayed 48 hr post transfection. The DNAs used were: FLAG-V2R, HA-ScFv control, HA-ScFv30, β-arrestin1-GFP. The antibodies used were: mouse anti-FLAG (M1, Sigma); rabbit anti-HA (cell signaling); goat anti-rabbit Alexa-Fluor647 and goat anti-mouse Alexa-Fluor555 (Thermo Fisher).

### Co-immunoprecipitation assay

In order to screen for Fabs that selectively recognized active conformation of βarr1, FLAG-tagged Receptor (3.5μg) along with βarr1 (3.5μg) in ratio 1:1 was overexpressed in HEK-293 cells using PEI based transient transfection. 48h post-transfection, cells were serum starved for at least 4 hrs, lysed by douncing and incubated with either Fab4,7,9 or 12 for 1h at room temperature to allow a stable Receptor-βarr1-Fab complex. Subsequently, 20 μl of pre-equilibrated Protein L (Capto L, GE Healthcare) beads (20mM HEPES, 150mM NaCl) was added to the mixture and allowed to incubate for additional 1h. Beads were washed thrice with wash buffer containing 20mM HEPES, 150mM NaCl supplemented with 0.01% MNG and eluted with 2X SDS loading buffer. Eluted samples were run on 12% SDS-polyacrylamide gel electrophoresis and probed for receptor using HRP-coupled anti-FLAG M2 antibody (Sigma, 1:2000). Fabs were visualized on gel using Coomassie staining.

In order to assess the ability of intrabodies (Ib30 and Ib4) with only activated form of βarr1, HEK-293 cells overexpressing FLAG-tagged Receptor (2.3μg), βarr1 (2.3μg) and HA-tagged Ibs (2.3μg) were stimulated with agonist at saturated concentration 48h post-transfection. Cells were lysed in NP-40 lysis buffer (Tris 50 mM; NaCl 150mM; PhosStop 1X; Protease inhibitor 1X; NP-40 1%) followed by incubation with 20μl of pre-equilibrated HA beads (Sigma, A-2095) for 2h at 4°C. Beads were washed thrice with wash buffer containing 20mM HEPES, 150mM NaCl maintaining cold conditions. For elution 2X SDS loading buffer was used. FLAG-tagged receptor was probed using HRP-coupled anti-FLAG M2 antibody (1:2000) while HA-tagged Ibs were detected using HA-probe antibody (sc-805, 1:5000). Ib-CTL which does not recognize β-arr1 was used as a negative control in these experiments. In order to further confirm that the Ib30 recognizes a functional Receptor-βarr1 complex, we overexpressed FLAG-tagged β2V2R, βarr1 and HA-tagged Ib30 in HEK-293 cells. 48h post-transfection cells were stimulated with agonists of varying efficacies (Inverse agonist: ICI and carazolol; partial agonists: salmeterol, salbutamol, clenbuterol and norepinephrine; full agonists: Isoproterenol) and co-immunoprecipitated using HA-beads agarose by the same method described above.

To determine the conformation of βarr1 bound to C5aR1-WT receptor versus C5aR1-V2 chimera, surface expression of the two receptors were first optimized by Surface ELISA and subsequently transfection was carried in a way that would yield almost equal surface expression for the two receptors. Accordingly, HEK-293 cells were transfected with βarr1 (3.5 μg) along with FLAG-tagged C5aR1-WT (3.5 μg) or FLAG-tagged C5aR1-V2 (2.5 μg). 48 hrs post-transfection cells were starved for 4h and stimulated with agonist (C5a, 100 nM). Cells were lysed by douncing and then further incubated with 5 μg of Fab30 for 1h at room temperature to enable Receptor-βarr1-Fab30 complex formation. Subsequently, 20 μl of pre-equilibrated Protein L (Capto L, GE Healthcare) beads (20mM HEPES, 150mM NaCl) was added to the mixture and allowed to incubate for additional 1h. Beads were washed thrice with wash buffer containing 20mM HEPES, 150mM NaCl supplemented with 0.01% MNG and eluted with 2× SDS loading buffer. Eluted samples were run on 12% SDS-polyacrylamide gel electrophoresis and probed for receptor using HRP-coupled anti-FLAG M2 antibody (Sigma, 1:2000). Fabs were visualized on gel using Coomassie staining. A separate gel containing 20 μl of the whole cell lysate was run on 12% gel and probed with anti-FLAG M2 antibody to compare expression of both WT and chimeric receptors.

In order to determine the interacting interface of the third intracellular loop (ICL3) in M2R with βarr1, purified βarr1 were pre-incubated with a 10-fold molar excess of V2Rpp or alone for 30 mins at room temperature. HEK-293 cell lysate overexpressing the M2R and stimulated with its agonist carbachol (20μM) at 0, 5 and 15 min time points were subsequently added to the βarr1 alone or βarr1 activated and saturated with V2Rpp. The reaction was allowed to proceed for 1hr at room temperature. Fab30 (2.5μg/reaction) was also kept in the reaction for stabilizing the M2R-βarr1 complex. Reactions with no Fab30 were kept as a negative control. Following an incubation of 1 hr, pre-equilibrated M1-Flag agarose beads (20uL) were added to the reaction mixture supplemented with 2mM CaCl_2_ and incubated for an additional 1h at room temperature. Beads were extensively washed twice with low salt wash buffer (20mM HEPES pH 7.4, 150mM NaCl, 2mM CaCl_2_, 0.01% MNG) first followed by washes in high salt buffer (20mM HEPES pH 7.4, 150mM NaCl, 2mM CaCl_2_, 0.01% MNG) with additional washes (twice) in low salt buffer again. Bound proteins were eluted using elution buffer containing FLAG peptide (20mM HEPES pH 7.4, 150mM NaCl, 2mM EDTA, 250 μg/ml FLAG peptide, 0.01% MNG,) and separated by running on 12 % SDS-PAGE and visualized by Western Blotting (βarr antibody, 1:5000; anti-FLAG M2 antibody, 1:5000) (Figure 4C).

### Confocal microscopy

In order to visualize the role of Ib30/Ib4 as a biosensor of GPCR activation co-localization of Ibs with activated βarr1 was monitored using confocal microscopy. HEK-293 cells co-transfected with Receptor (2.3μg), βarr1-m-cherry (2.3μg) and Ib-YFP (2.3μg) each. After 24 h, cells were seeded on to cell culture treated confocal dishes (GenetiX; 100350) already coated with 0.01% poly-D-lysine (Sigma). 48h post-transfection cells were serum starved for at least 6h prior to stimulation with indicated agonists at saturating concentrations. For confocal live imaging we used Zeiss LSM 710 NLO confocal microscope and samples were housed on a motorized XY stage with a CO2 enclosure and a temperature controlled platform equipped with 32x array GaAsP descanned detector (Zeiss). YFP was excited with a diode laser at 488 nm laser line while m-cherry was excited at 561 nm. Laser intensity and pinhole settings were kept in the same range for parallel set of experiments and spectral overlap for any two channels was avoided by adjusting proper filter excitation regions and bandwidths. Images were scanned using the line scan mode and images were finally processed in ZEN lite (ZEN-blue/ZEN-black) software suite from ZEISS. Line-scan analysis was performed using ImageJ plot profile plug-in to measure fluorescence intensities across a drawn line. Graphs were plotted after intensities were normalized by subtracting background.

### ERK1/2 MAP kinase phosphorylation assay

For assessing the role of βarr1-mediated ERK2 activation on V2R, AT1AR, B_2_R, B_2_R^ΔG/L370T^ and B_2_R^L370T^, HEK-293 cells with selective knockdown of βarr1, attained by sh-RNA mediated transfections were used. shRNA mediated knockdown reduced the amount of βarr1 by ∼50%. Stable selection was maintained by inclusion of G418 (1.5μg/ml) in the growth medium. About 3 million cells were seeded on a 10 cm cell-culture dish a day prior to transfection to achieve ∼ 60% confluency. For transfections, 0.5μg, 1μg and 2μg of V2R, B2R and AT1AR were used respectively. After 24 hrs, 1 million cells were split on to a 6 well plate. 48h post-transfection cells were serum starved for at least 5 hrs in serum-free medium supplemented with 10mM HEPES (pH 7.4) and 0.1% Bovine serum albumin. Cells were stimulated with an increasing dose of agonist (-log9, -log8 and –log7) and were lysed using 2X SDS loading buffer, boiled at 95°C for 15 mins.

In order to examine the effect of Ib30 on βarr1 mediated ERK activation, HEK-293 cells were co-transfected with human Vasopressin receptor (V2R; 3.5μg) and either Ib30 or Ib-CTL (3.5μg) using PEI based transient transfection. After 24h, one million cells were plated on a six-well plate. 48h post-transfection cells were serum starved for at least 4h and stimulated with agonist (AVP,100 nM). Following stimulation cells were lysed using 2X SDS loading buffer, boiled at 95°C for 15 mins and loaded onto 12% SDS-polyacrylamide gel electrophoresis. All experiments were carried out on HEK-293 cells with low passage and maintained in Dulbecco’s modified Eagle’s complete media (Sigma) supplemented with 10% fetal bovine serum (Thermo Scientific) and 1% penicillin–streptomycin at 37⍰°C under 5% CO2.

For detecting phosphorylation of ERK1/2 in both HEK-293 cells and knockdown cells Western blotting analysis of the whole-cell lysate was performed and transferred to polyvinylidene difluoride membranes (PVDF;BioRad). The membrane was blocked with 5% BSA (SRL) for 11h and then probed with anti-pERK primary antibody (CST, catalog number. 9101; 1:5,000 dilution) overnight at 4⍰°C followed by 11h incubation with anti-rabbit IgG secondary antibody (Genscript, catalog number. A00098) at room temperature. The membrane was then washed with 1 × TBST thrice and developed using Chemi Doc (BioRad). The anti-pERK antibody was stripped-off using 1× stripping buffer and then reprobed with anti-tERK antibody (CST, catalog number. 9102 and 4695; 1:5,000 dilution).

### Three-channel confocal Imaging Imaging

Receptor imaging of live or fixed cells was monitored by “feeding” cells with Alexa-Fluor555-conjugated FLAG antibody (15 min, 37°C) in phenol-red-free DMEM prior to agonist treatment. Fixed cells were washed three times in PBS/0.04% EDTA to remove FLAG antibody bound to the remaining surface receptors, fixed using 4% PFA (20 min at RT), permeabilized and stained using HA primary antibody followed by Alexa-Fluor647 secondary antibody. For co-localization of FLAG-V2R with endosomal markers, cells were treated as above except incubated with either of the following primary antibodies post-permeabilization; EEA1 (rabbit anti-EEA1 antibody from Cell Signaling Technology) or APPL1 (rabbit anti-APPL1 antibody from Cell Signaling Technology). Cells were imaged using a TCS-SP5 confocal microscope (Leica) with a 63× 1.4 numerical aperture (NA) objective and solid-state lasers of 488 nm, 561 nm, and/or 642 nm as light sources. Leica LAS AF image acquisition software was utilized. All subsequent raw-image tiff files were analyzed using ImageJ or LAS AF Lite (Leica).

### GloSensor Assay

To assess the role of Ib30 on G protein signalling and desensitization GloSensor assay was performed with only vasopressin receptor system. HEK-293 cells were triple transfected with human Vasopressin receptor (V2R receptor; 2.3μg), the luciferase-based cAMP biosensor (2.3 μg; pGloSensorTM-22F plasmid; Promega) and Ib30/Ib-CTL (2.3 μg) using PEI based transient transfections. 14–16 h post-transfection, media was aspirated and cells were flushed and pooled together in assay buffer containing 1× Hanks balanced salt solution, pH 7.4 and 20 mM of 4-(2-hydroxyethyl)-1-piperazineethanesulfonic acid [HEPES]. Density of the cells was determined by counting on haemocytometer and subsequently the volume of cells that should yield 125,000 cells per 100μl in a 96 well plate was calculated. Cells were pelleted at 2000 rpm for 3 mins to remove the assay buffer and the pellet was resuspended in the desired volume of sodium luciferin solution prepared in the same assay buffer. After seeding the cells in a 96 well plate it was allowed to incubate at 37°C for 90 min followed by an additional incubation of 30 mins at room temperature. For stimulation various doses of ligand (AVP) were prepared by serial dilution ranging from 0.1 pM to 1 μM and added to the cells with help of a multichannel pipette. Luminescence was recorded using a microplate reader (Victor X4; Perkin Elmer). Effect of Ib30 was compared to Ib-CTL and normalized with respect to maximal stimulation by agonist (treated as 100%). Data were plotted and analyzed using nonlinear regression in GraphPad Prism software.

### BRET assay

Transient transfections were performed on cells seeded (40,000 cells/100 µl/well) in white 96-well microplates (Greiner) using 25 kDa linear polyethylenimine (PEI) as transfecting agent, at a ratio of 4:1 PEI/DNA. To monitor receptor or β-arrestin trafficking from the cell surface or in the endosomes, we used enhanced bystander BRET (ebBRET) where the BRET acceptor, a *Renilla* green fluorescent protein (rGFP), is fused to either CAAX or FYVE domains from Kras and endofin proteins respectively to target respectively the plasma membrane (rGFP-CAAX) or the early endosomes (rGFP-FYVE); the receptor or β-arrestin are fused to the BRET donor *Renilla* luciferase II (RlucII) (*22*). To monitor β-arrestin interaction with V2R, we used β-arrestin fused to RlucII and V2R fused to the yellow variant of the *aquaria Victoria* green fluorescent protein, YFP (*23*). Forty-eight hours later, culture media was removed, cells were washed with DPBS (Dulbecco’s Phosphate Buffered Saline) and replaced by HBSS (Hank’s Balanced Salt Solution). For time-course experiment, after a 3 minutes pre-incubation with of 2.5 µM coelenterazine H (BRET1 for V2R β-arrestin interaction) from Goldbio or coelenterazine 400a (BRET2 for CAAX and FYVE assays) from Nanolight Technology, cells were stimulated with vehicle or AVP (100 nM) and BRET was measured every 45 seconds for 20 minutes. For concentration-response curves, cells were stimulated with increasing concentrations of AVP for 10 minutes and 2.5µM coelenterazine H (BRET1) or coelenterazine 400a (BRET2) was added 5 minutes before BRET measurement. BRET signals were recorded on a Mithras (Berthold scientific) microplate reader equipped with the following filters: 480/20⍰nm (donor) and 530/20⍰nm (acceptor) for BRET1 or 400/70⍰nm (donor) and 515/20⍰nm (acceptor) for BRET2. The BRET signal was determined as the ratio of the light emitted by the energy acceptor over the light emitted by energy donor. The agonist-promoted BRET signal (ΔBRET) was obtained by subtracting the BRET signal recorded in the presence of vehicle from that obtained following AVP treatment.

### Receptor and intrabody expression

To assess the receptor and intrabodies expression levels, we took advantage of the FLAG epitope fused at the N-terminus of the V2R (FLAG-V2R) and a HA epitope tag fused at the C-terminus of the intrabodies. Their relative expression was monitored by ELISA using anti-FLAG-HRP (Sigma) or anti-HA-HRP antibodies (Roche Diagnostics). Cells were fixed with 3% formaldehyde for 10 min and permeabilized with 0.1% triton X-100 to monitor the intrabodies expression. Cells were washed three times in washing buffer (1% BSA in DPBS), followed by a blocking step by incubating the cells for 1 hour in washing buffer. The horse radish peroxidase-conjugated antibody was then added for 1h at room temperature and the HRP activity was measured by adding o-phenylenediamine dihydrochloride (Sigma–Aldrich). The reaction was stopped by adding 0.6 M HCl followed by absorbance measurement at 492nm using a SpectraMax 190 plate reader (Molecular Devices).

### Molecular Dynamics simulation

In order to generate the complexes of the B2Rpp WT, the L370pT mutant of B2Rpp and the V2Rpp bound to βarr1, we used the structure of V2Rpp in complex with βarr1 (PDB code: 4JQI). Missing fragments in the βarr1 and V2Rpp structures were modelled using the loop modeler module available in the MOE package (https://www.chemcomp.com). B2Rpp and B2Rpp mutant systems were obtained by converting the sequence of the V2Rpp into that of the B2Rpp and the corresponding L370pT mutant. Finally, the co-crystallized FAB30 antibody was removed.

The complexes were solvated in TIP3P water, with the ionic strength kept at 0.15 M using NaCl ions. Simulation parameters were obtained from the Charmm36M forcefield1. Systems generated this way were simulated using the ACEMD software package2. To allow rearrangement of waters and sidechains, we carried out a 25ns equilibration phase in NPT conditions with restraints applied to backbone atoms. The timestep was set to 2 fs and the pressure was kept constant, using the Berendsen barostat. After the equilibration, systems were simulated in NVT conditions for 1μs in 4 parallel runs employing a 4fs timestep. For all runs temperature was kept at 300 K using the Langevin thermostat and hydrogen bonds were restrained using the RATTLE algorithm. Non-bonded interactions were cut-off at 9 Å with a smooth switching function applied at 7.5 Å. Before carrying out the structural analysis, simulation frames were aligned using the backbone atoms of the βarr1. To assess the magnitude of salt bridge formation between phosphorylated threonines (pT) residues and K294, we quantified frames in which the protonated nitrogen of K294 and oxygens of the phosphate group of each respective pT adopted a distance of 3.2 Å or less. Conformational variability of the lariat loop was studied with the clustering tool available in VMD3. As a clustering parameter we used RMSD (cutoff: 2.2) of the backbone atoms of residues 293 to 297 within the lariat loop.

