## Supplemental Figures for "Genetically encoded intrabody sensors illuminate structural and functional diversity in GPCR-β-arrestin complexes"

**Supplementary Figure 1.**

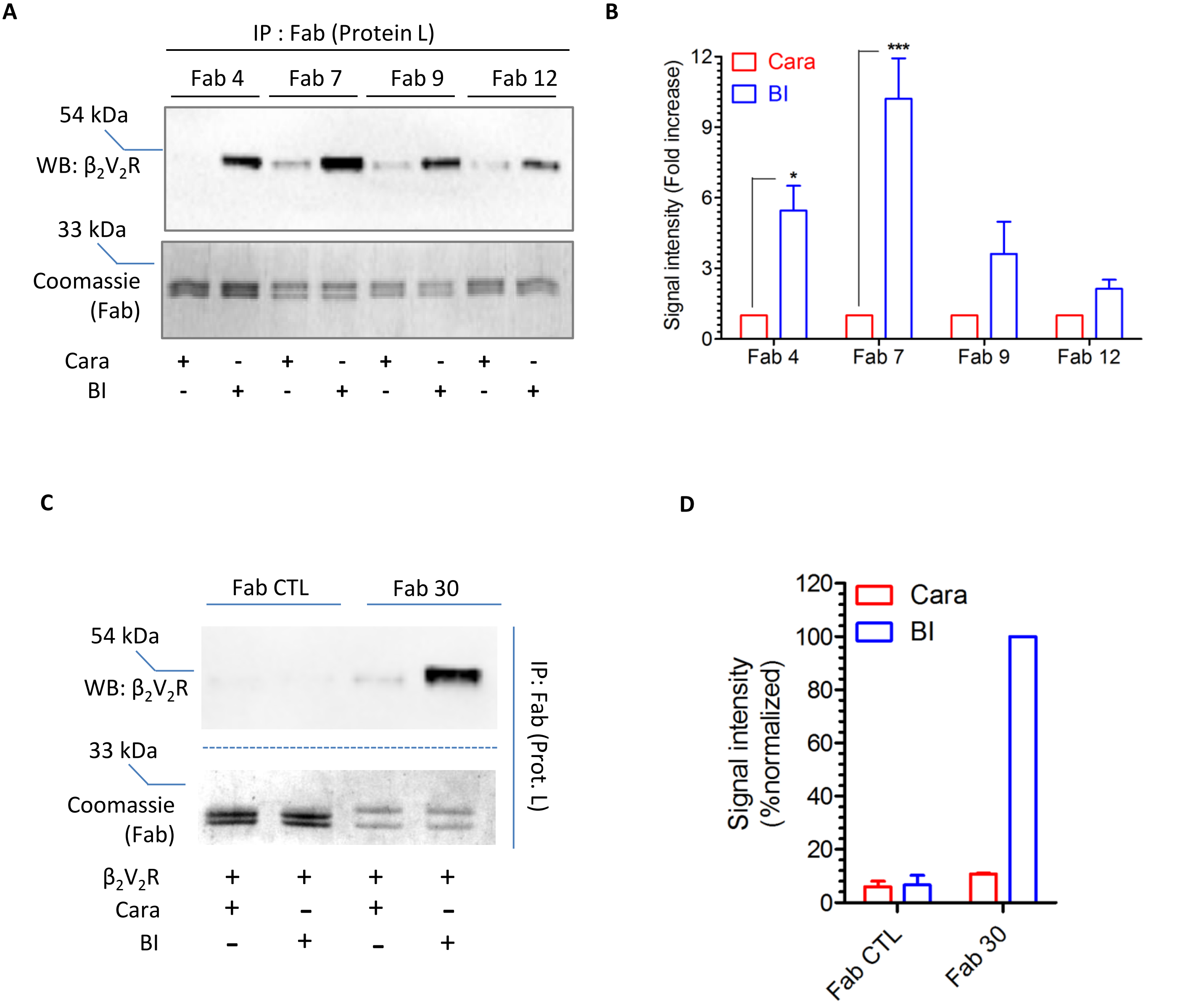


**Supplementary Figure.1. Screening and characterization of Fabs that selectively recognize Receptor-βarr1 complex *in vitro.*** **A** Synthetic antibody fragments (Fabs) that selectively recognize $\beta_2V_2R$-βarr1 complex. Cellular lysate from Sf9 cells, overexpressing the FLAG-tagged $\beta_2V_2R$ were mixed with purified βarr1 andFabs, stimulated with either an inverse-agonist (Carazolol 1 µM) or agonist (BI-167107 100 nM) followed by co-immunoprecipitation to allow complex formation of Receptor-βarr1-Fab. The complex was visualized by immobilizing the Fabs on Protein L beads and probing for $\beta_2V_2R$ using anti-FLAG M2 antibody byimmunoblotting. **B** Ability of Fabs to form stable $\beta_2V_2R$-βarr1 complex as obtained in A is shown as bar graph. Data shows densitometry based quantification andrepresents mean ± s.e.m of five independent experiments and is expressed as fold change of Fab stabilized complex formation in presence of an inverse agonist treated as one. **C** HEK-293 cells expressing $\beta_2V_2R$, βarr1 and Fab30 (or Fab-CTL) were stimulated with either an inverse-agonist (Carazolol 1 µM) or agonist (BI-167107 100 nM) followed by co-immunoprecipitation using Protein L beads as described in panel A. **D** The ability of Fab30 to recognize agonist-activated $\beta_2V_2R$-βarr1complex as shown in panel C is represented as bar graph. Data shows densitometry based quantification and represents mean ± s.e.m of two independent experiments and has been normalized by treating $\beta_2V_2R$-βarr1-Fab30 complex formation in presence of an agonist as 100%.

**Supplementary Figure 2.**

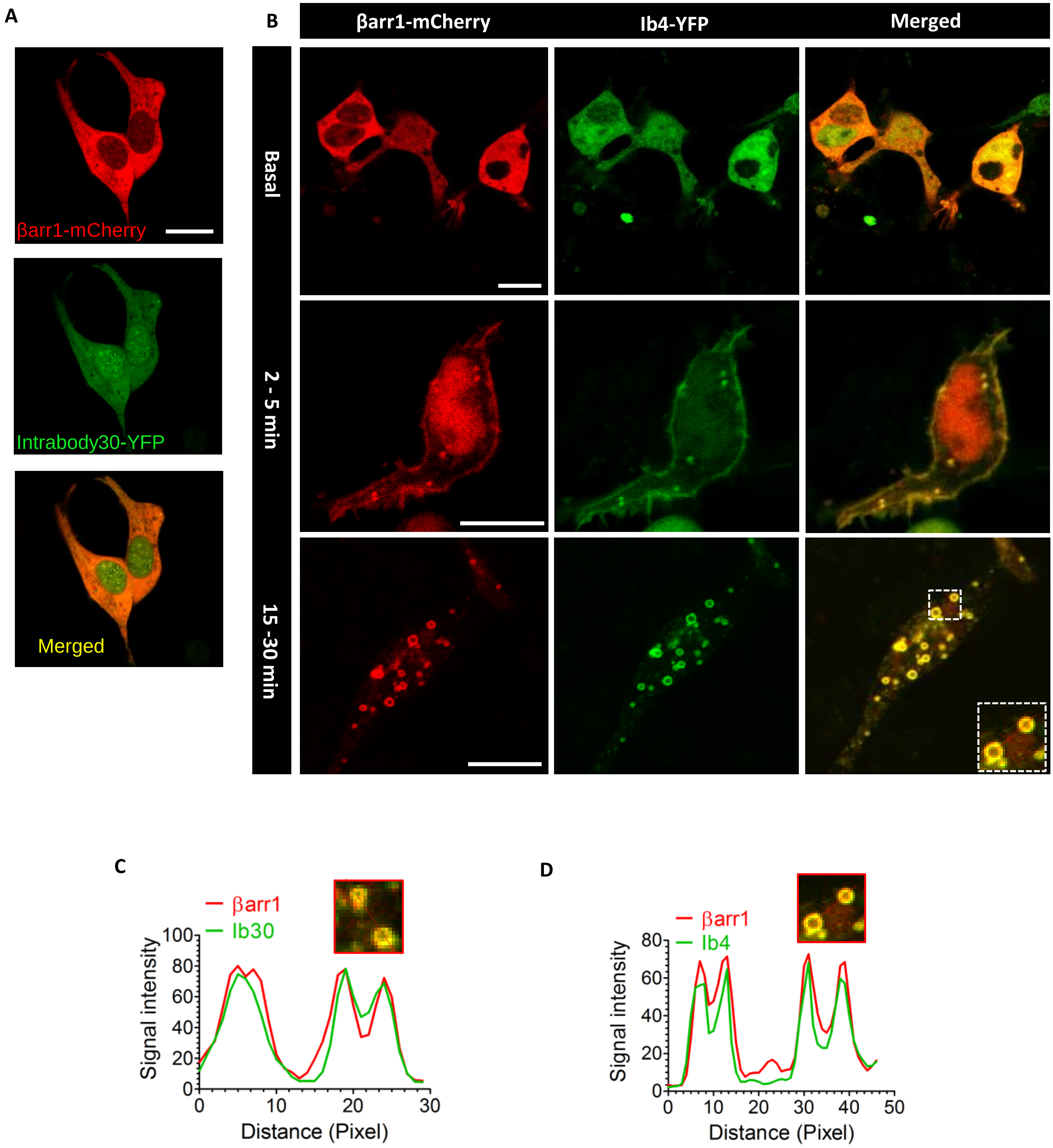


**Supplementary Figure.2. Intrabody4 reports agonist-induced βarr1 trafficking with spatio-temporal resolution.** HEK-293 cells overexpressing the $\beta_2V_2R$, βarr1-mCherry and YFP-tagged Ib4 were stimulated with the agonist (Isoproterenol 10 μM) at the indicated time points and localization of both βarr1 and Ib4 were monitored using live-cell imaging by confocal microscopy. Inset shows line scan analysis of fluorescence intensities from both channels (βarr1-mCherry and YFP-Ib4).Overlap of both fluorescence intensities suggests co-localization of Ib4 with activated βarr1. Scale bar is 10 μM.

**Supplementary Figure 3.**

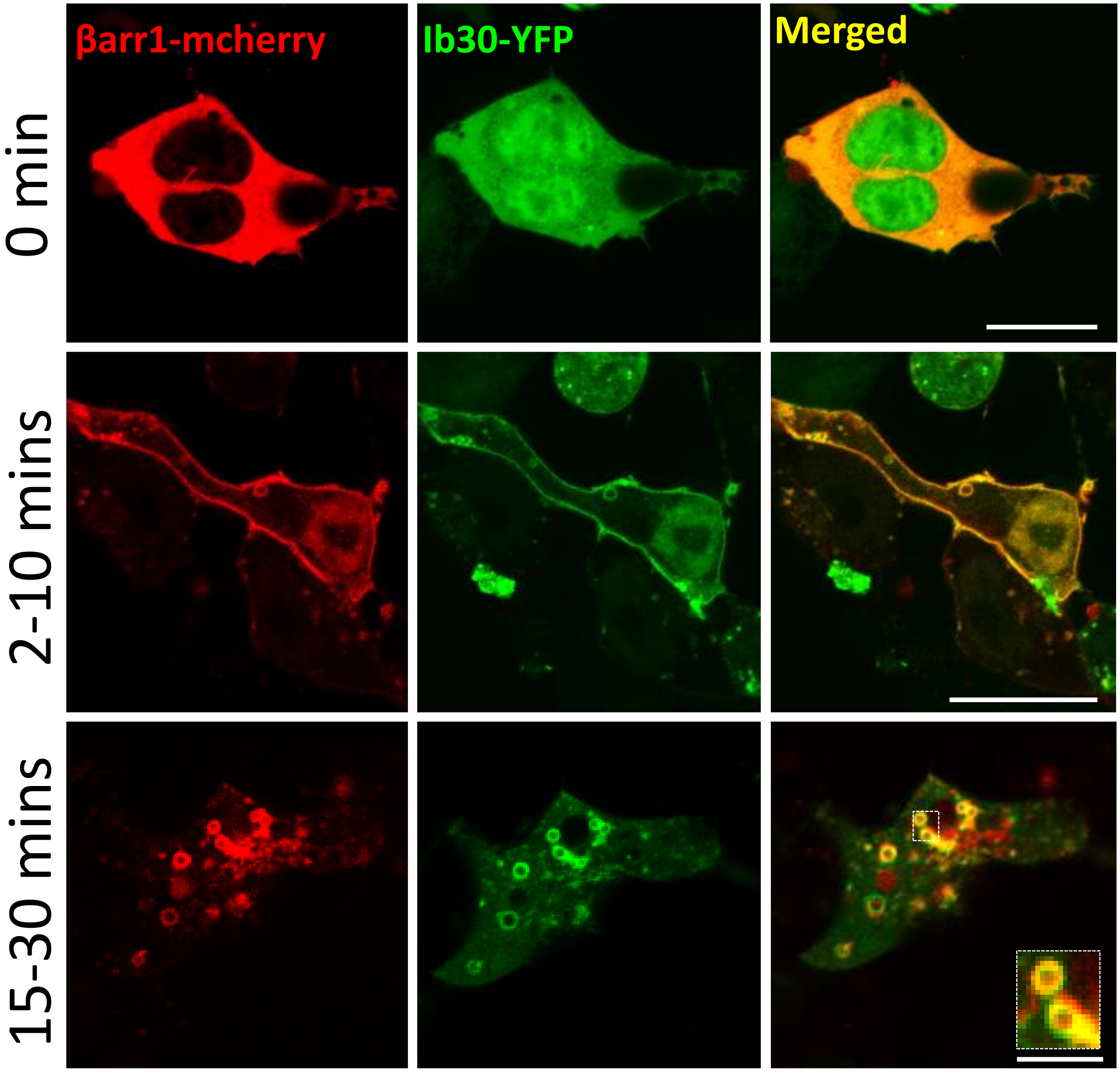


**Supplementary Figure 3. Intrabody30 sensor reports βarr1 recruitment and trafficking for the α2B-V2R.** HEK-293 cells were transfected with α2B-V2R, βarrestin1-mCherry and YFP-tagged Ib30, and agonist-induced receptor trafficking was assessed using confocal microscopy. Under unstimulated conditions (0 min) both βarr1-mCherry and Ib30-YFP shows cytoplasmic distribution. Within 2-10 mins of agonist stimulation (epinephrine 20 μM,) both βarr1-mCherry and Ib30-YFP are localized to the plasma membrane. Upon prolonged agonist-exposure, (15-30 mins), Ib30-YFP colocalizes with βarr1-mCherry in endosomal vesicles as apparent by the appearance of doughnut like structures. Scale bar is 10μm.

**Supplementary Figure 4.**

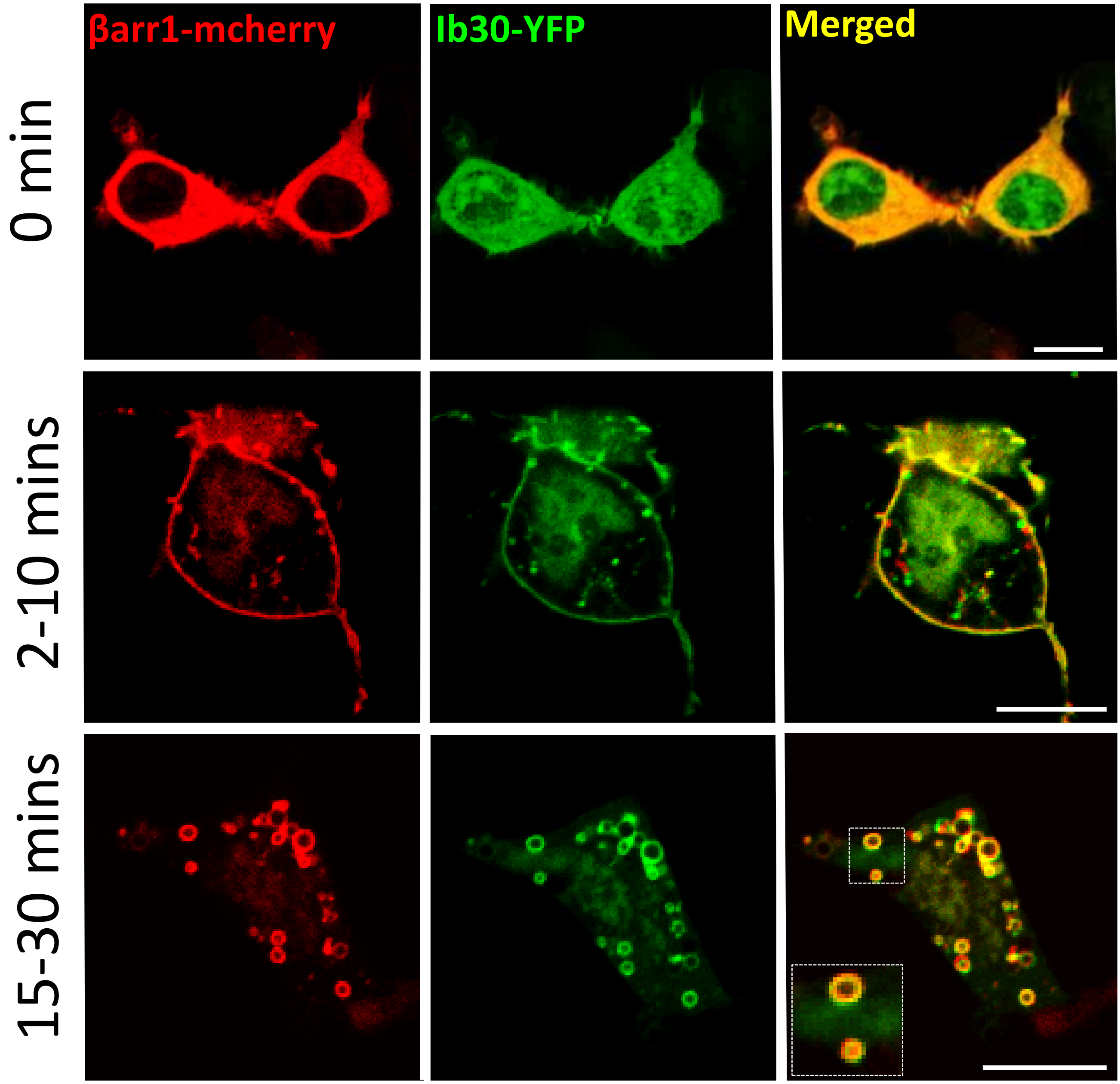


**Supplementary Figure 4. Intrabody30 sensor reports βarr1 recruitment and trafficking for the CCR2-V2R.** HEK-293 cells were transfected with -V2R, βarrestin1-mCherry and YFP-tagged Ib30, and agonist-induced receptor trafficking was assessed using confocal microscopy. Under unstimulated conditions (0 min) both βarr1-mCherry and Ib30-YFP shows cytoplasmic distribution. Within 2-10 mins of agonist stimulation (epinephrine 20 μM,) both βarr1-mCherry and Ib30-YFP are localized to the plasma membrane. Upon prolonged agonist-exposure, (15-30 mins), Ib30-YFP colocalizes with βarr1-mCherry in endosomal vesicles as apparent by the appearance of doughnut like structures. Scale bar is 10μm.

**Supplementary Figure 5.**

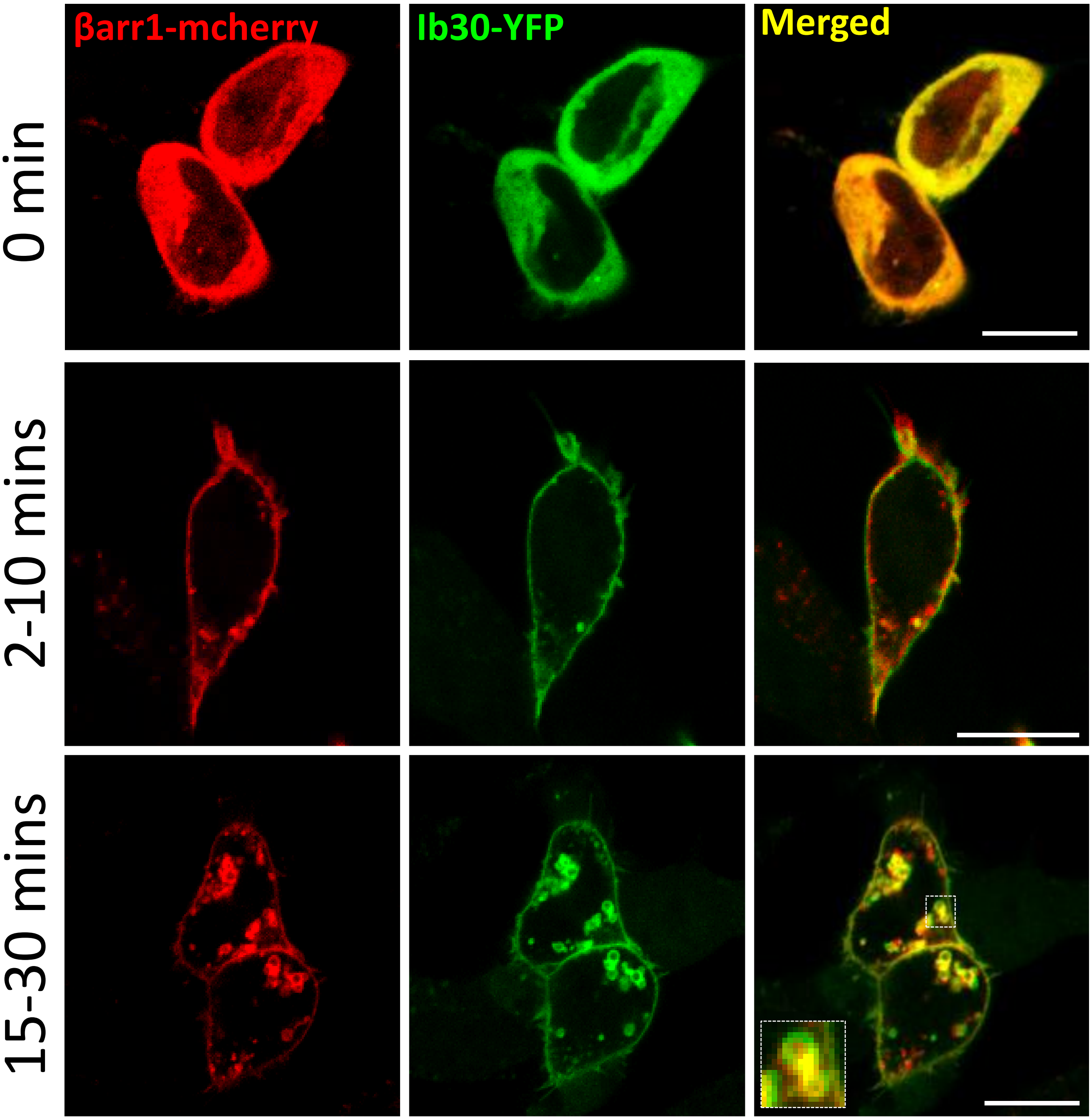


**Supplementary Figure 5. Intrabody30 sensor reports βarr1 recruitment and trafficking for the D2-V2R.** HEK-293 cells were transfected with D2-V2R, βarrestin1-mCherry and YFP-tagged Ib30, and agonist-induced receptor trafficking was assessed using confocal microscopy. Under unstimulated conditions (0 min) both βarr1-mCherry and Ib30-YFP shows cytoplasmic distribution. Within 2-10 mins of agonist stimulation (epinephrine 20 μM,) both βarr1-mCherry and Ib30-YFP are localized to the plasma membrane. Upon prolonged agonist-exposure, (15-30 mins), Ib30-YFP colocalizes with βarr1-mCherry in endosomal vesicles as apparent by the appearance of doughnut like structures. Scale bar is 10μm.

**Supplementary Figure 6.**

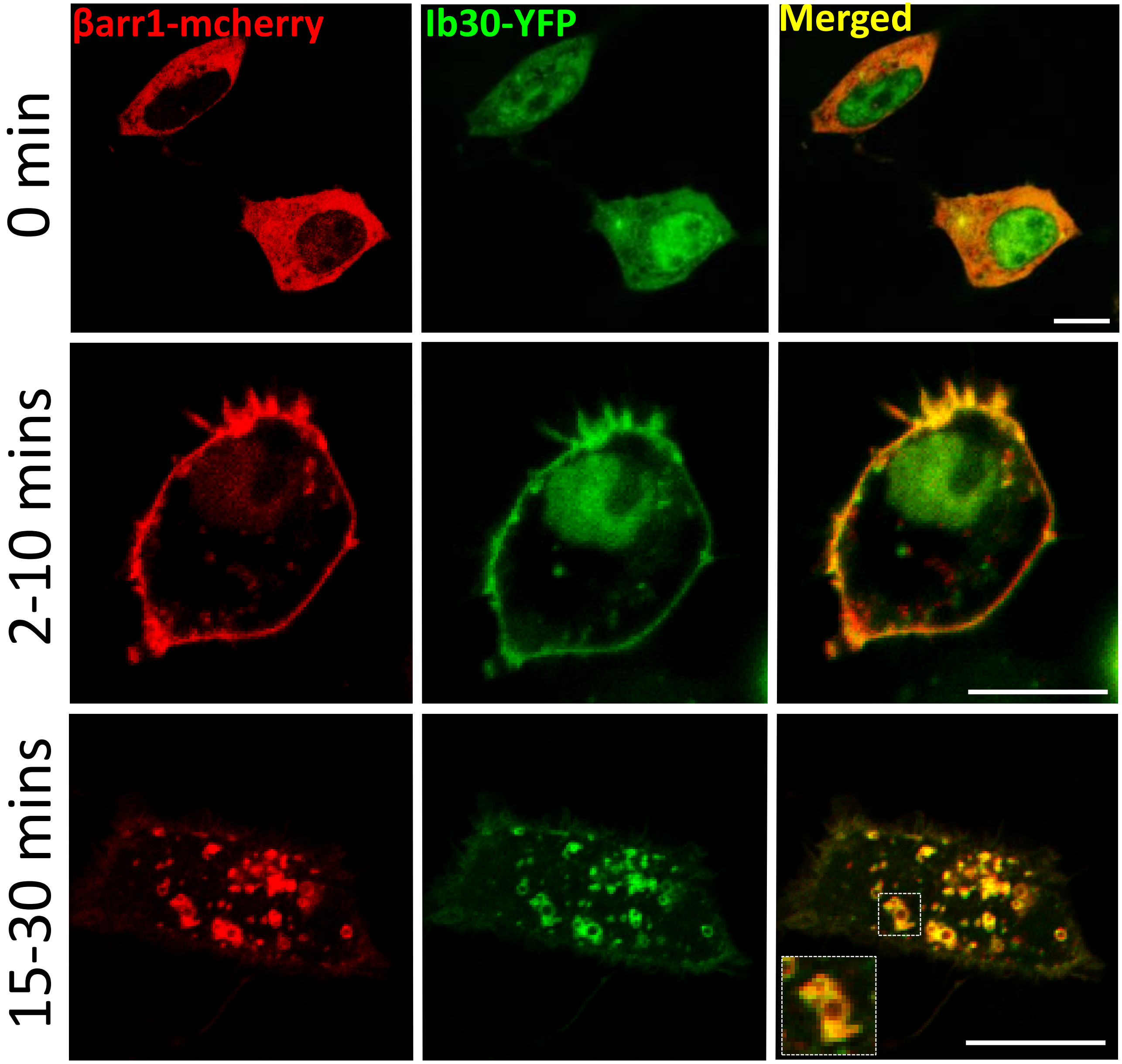


**Supplementary Figure 6. Intrabody30 sensor reports βarr1 recruitment and trafficking for the D5-V2R.** HEK-293 cells were transfected with D5-V2R, βarrestin1-mCherry and YFP-tagged Ib30, and agonist-induced receptor trafficking was assessed using confocal microscopy. Under unstimulated conditions (0 min) both βarr1-mCherry and Ib30-YFP shows cytoplasmic distribution. Within 2-10 mins of agonist stimulation (epinephrine 20 µM,) both βarr1-mCherry and Ib30-YFP are localized to the plasma membrane. Upon prolonged agonist-exposure, (15-30 mins), Ib30-YFP colocalizes with βarr1-mCherry in endosomal vesicles as apparent by the appearance of doughnut like structures. Scale bar is 10µm.

**Supplementary Figure 7.**

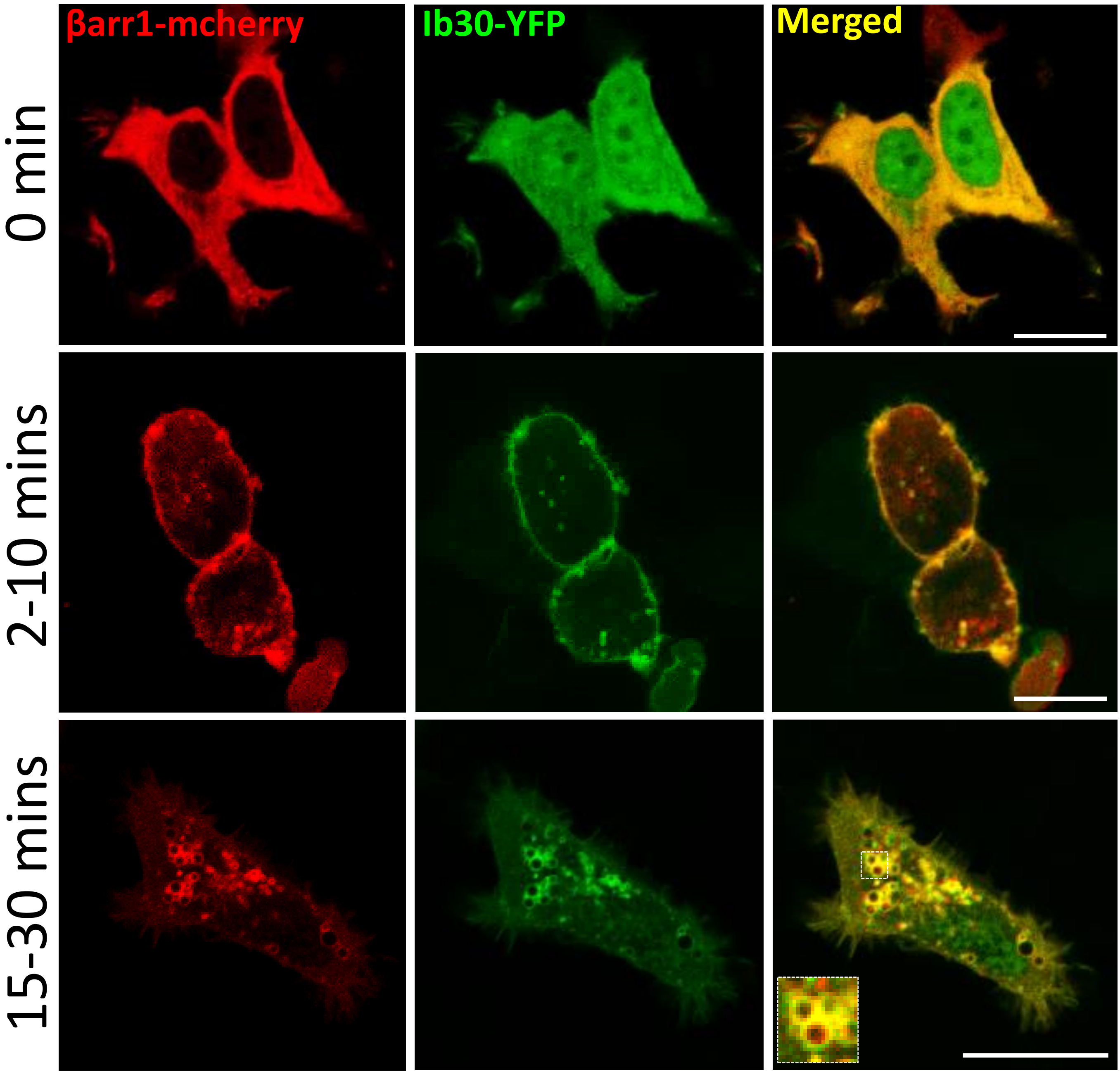


**Supplementary Figure 7. Intrabody30 sensor reports βarr1 recruitment and trafficking for the M5-V2R.** HEK-293 cells were transfected with M5-V2R, βarrestin1-mCherry and YFP-tagged Ib30, and agonist-induced receptor trafficking was assessed using confocal microscopy. Under unstimulated conditions (0 min) both βarr1-mCherry and Ib30-YFP shows cytoplasmic distribution. Within 2-10 mins of agonist stimulation (epinephrine 20 µM,) both βarr1-mCherry and Ib30-YFP are localized to the plasma membrane. Upon prolonged agonist-exposure, (15-30 mins), Ib30-YFP colocalizes with βarr1-mCherry in endosomal vesicles as apparent by the appearance of doughnut like structures. Scale bar is 10µm.

**Supplementary Figure 8.**

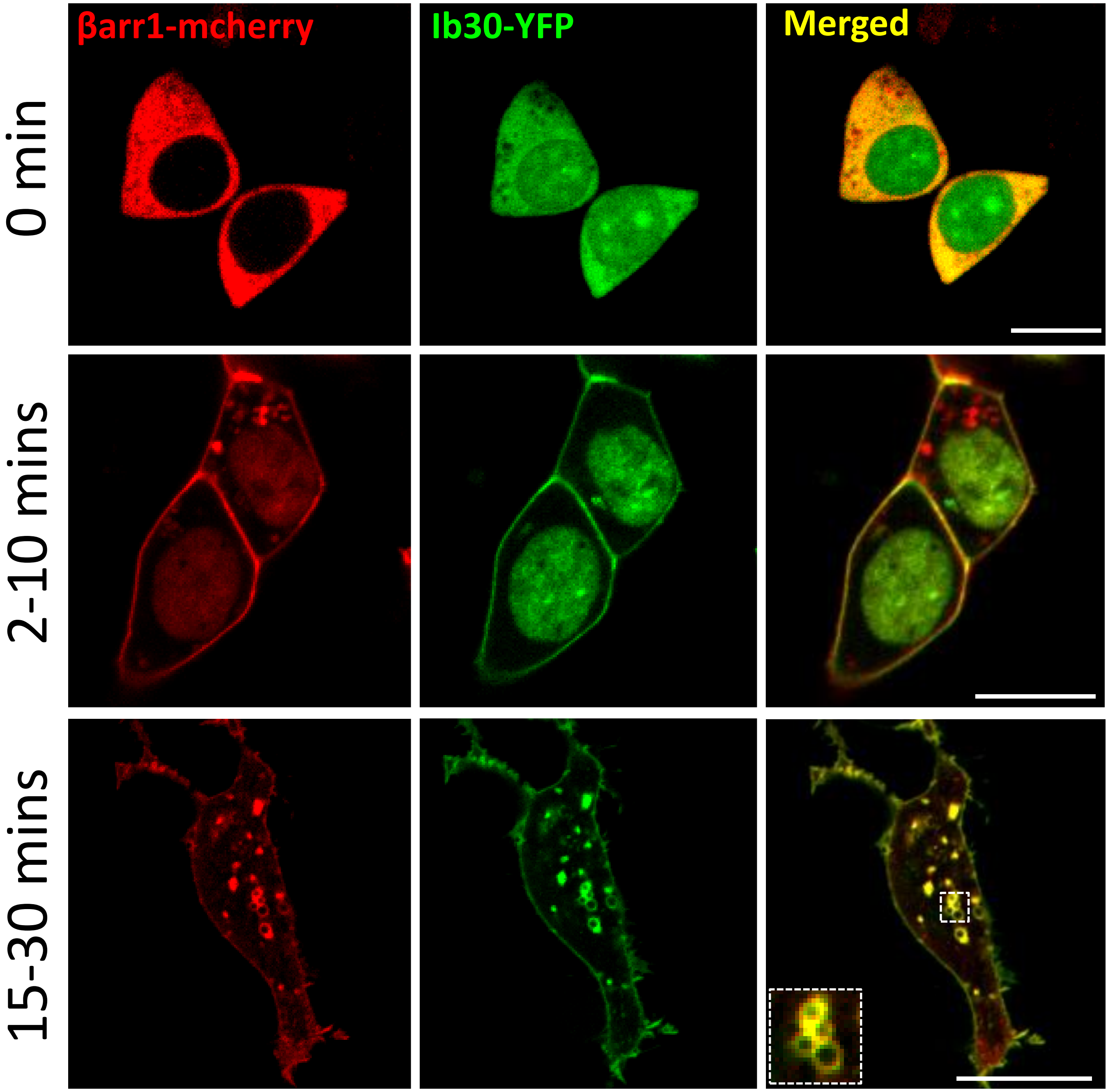


**Supplementary Figure 8. Intrabody30 sensor reports βarr1 recruitment and trafficking for the C5aR1-V2R.** HEK-293 cells were transfected with C5aR1-V2R, βarrestin1-mCherry and YFP-tagged Ib30, and agonist-induced receptor trafficking was assessed using confocal microscopy. Under unstimulated conditions (0 min) both βarr1-mCherry and Ib30-YFP shows cytoplasmic distribution. Within 2-10 mins of agonist stimulation (epinephrine 20 μM,) both βarr1-mCherry and Ib30-YFP are localized to the plasma membrane. Upon prolonged agonist-exposure, (15-30 mins), Ib30-YFP colocalizes with βarr1-mCherry in endosomal vesicles as apparent by the appearance of doughnut like structures. Scale bar is 10μm.

**Supplementary Figure 9.**

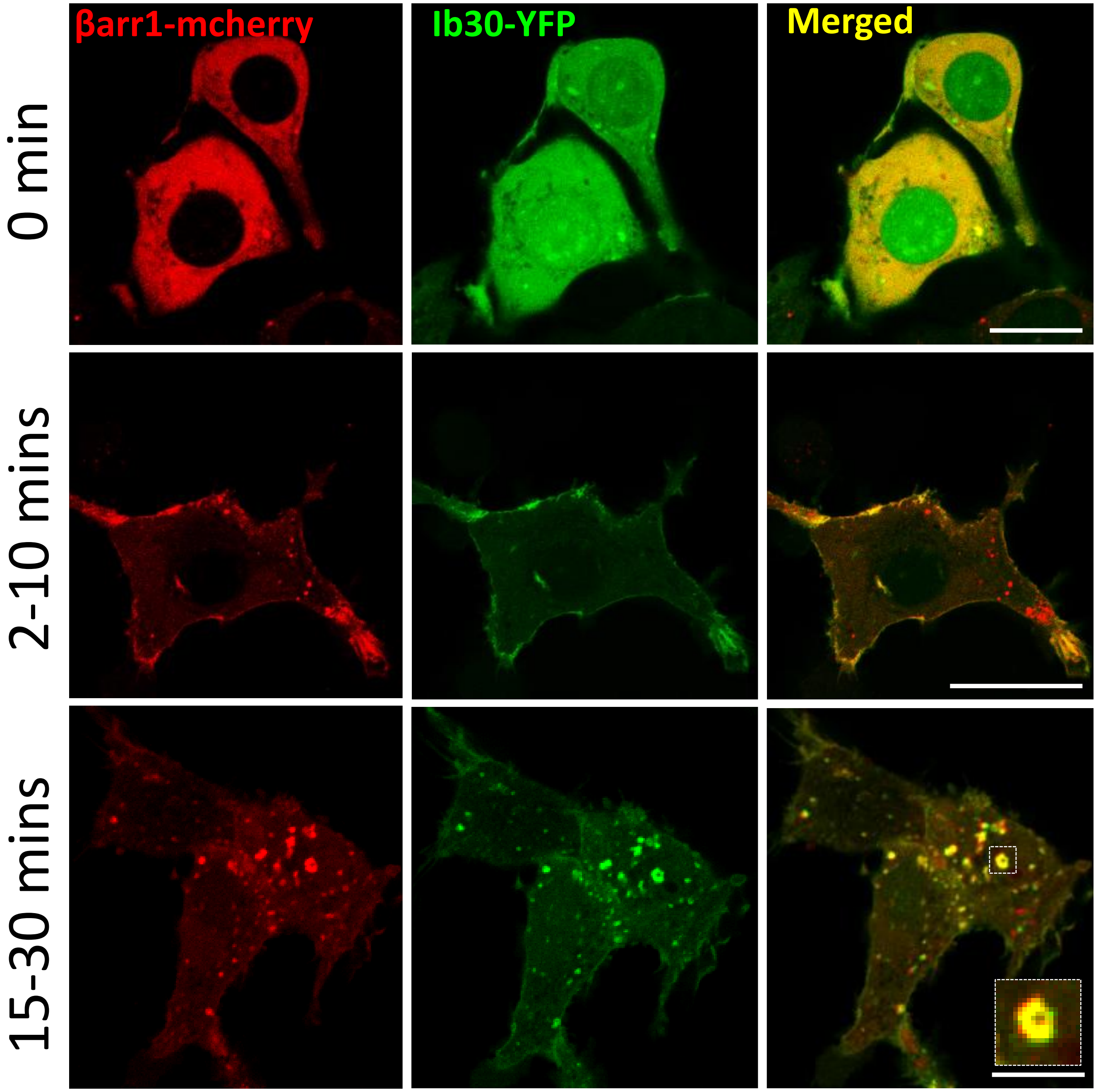


**Supplementary Figure 9. Intrabody30 sensor reports βarr1 recruitment and trafficking for the ACKR2-V2R.** HEK-293 cells were transfected with ACKR2-V2R, βarrestin1-mCherry and YFP-tagged Ib30, and agonist-induced receptor trafficking was assessed using confocal microscopy. Under unstimulated conditions (0 min) both βarr1-mCherry and Ib30-YFP shows cytoplasmic distribution. Within 2-10 mins of agonist stimulation (epinephrine 20 μM,) both βarr1-mCherry and Ib30-YFP are localized to the plasma membrane. Upon prolonged agonist-exposure, (15-30 mins), Ib30-YFP colocalizes with βarr1-mCherry in endosomal vesicles as apparent by the appearance of doughnut like structures. Scale bar is 10μm.

**Supplementary Figure 10.**

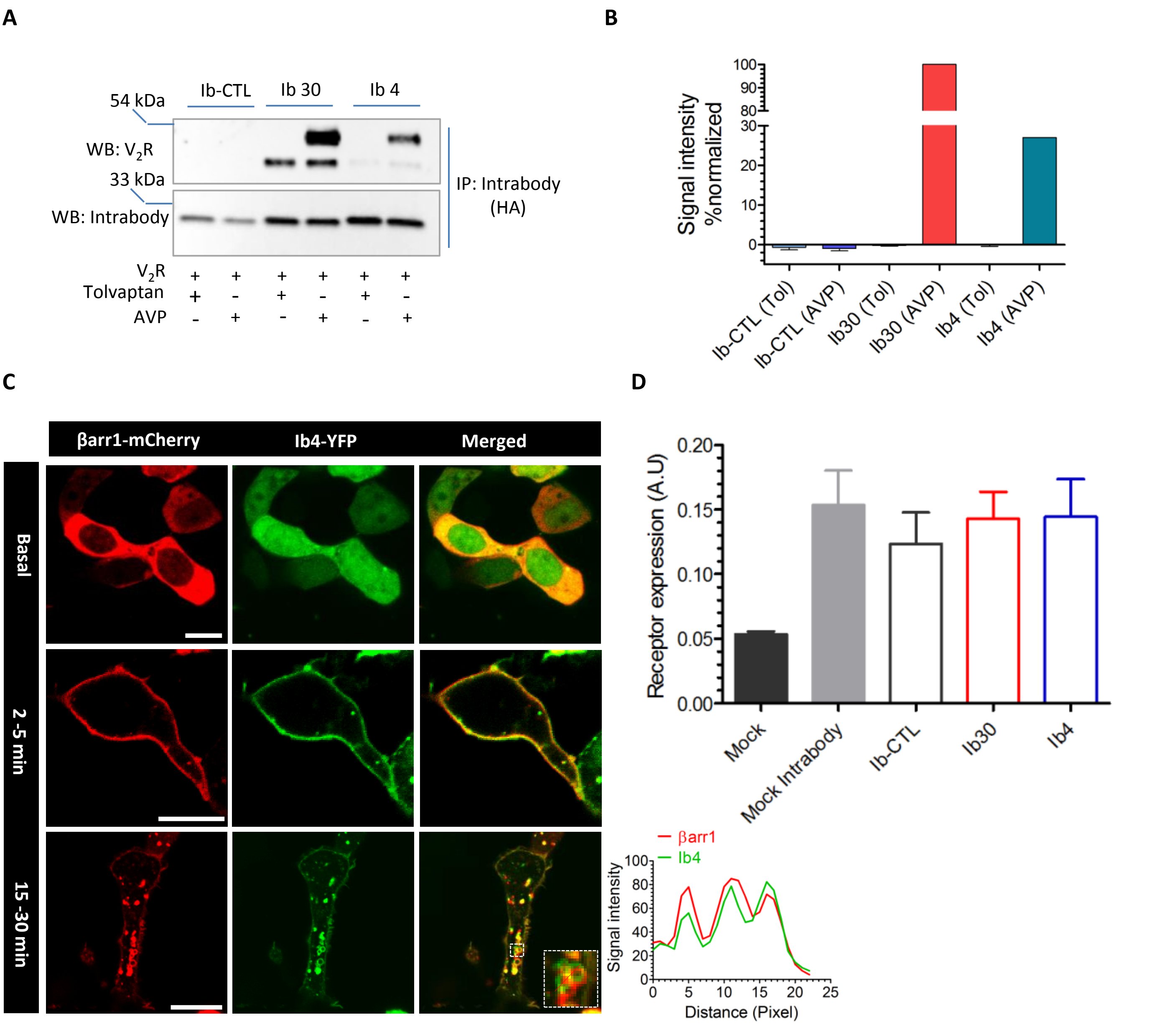


**Supplementary Figure 10. Intrabodies also recognize V2R-βarr1 complex with spatio-temporal resolution. A.** HEK-293 cells expressing V2R, βarr1 and intrabodies were stimulated with either an inverse-agonist (Tolvaptan) or agonist (AVP) followed by co-immunoprecipitation using anti-HA antibody agarose. The proteins were visualized by Western blotting using anti-Flag M2 antibody and anti-HA antibody. **B.** Densitometry-based quantification of the data presented in panel A (mean±s.e.m) from three independent experiments. **C.** Intrabody4 reports agonist-induced βarr1 trafficking. HEK-293 cells expressing V2R, βarr1-mCherry and Ib4-YFP were stimulated with agonist (AVP 100 nM) for indicated time-points and the localization of βarr1 and Ib4 were visualized using confocal microscopy. Scale bar is 10µm. **D.** Expression levels of Intrabodies were assessed by whole cell based Surface ELISA. HEK-293 cells were transfected with HA-tagged intrabodies and the relative expression was monitored 48h post-transfection using anti-HA-HRP antibodies

**Supplementary Figure 11.**

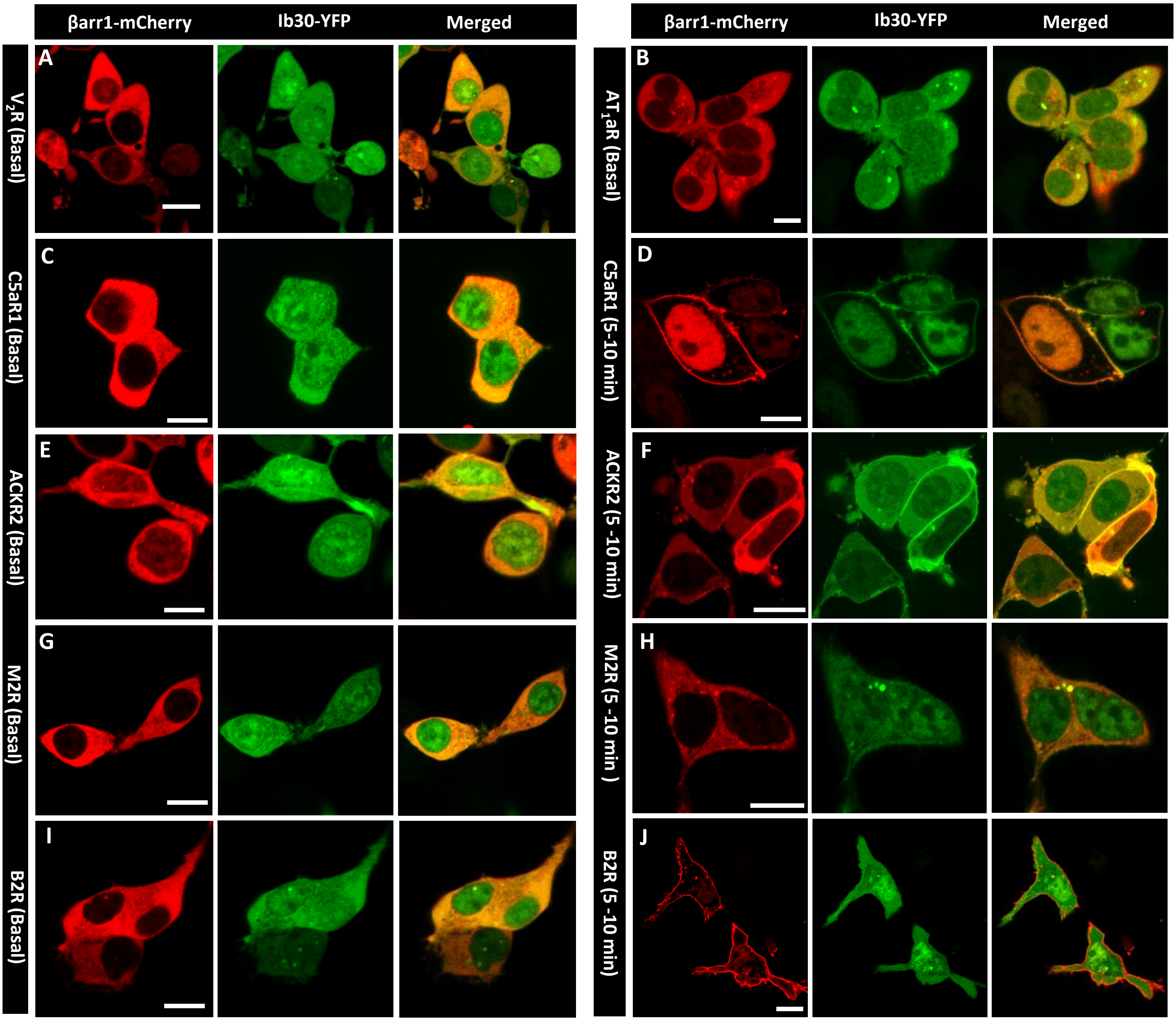


**Supplementary Figure 11. Intrabody30 reports recruitment and trafficking of βarr1 for broad set of native Class B GPCRs.** HEK-293 cells expressing the respective receptor, βarr1-mCherry and Ib30-YFP were stimulated with agonist for indicated time-points and the localization of βarr1 and Ib30 were visualized using confocal microscopy. Scale bar is 10µm.

**Supplementary Figure 12.**

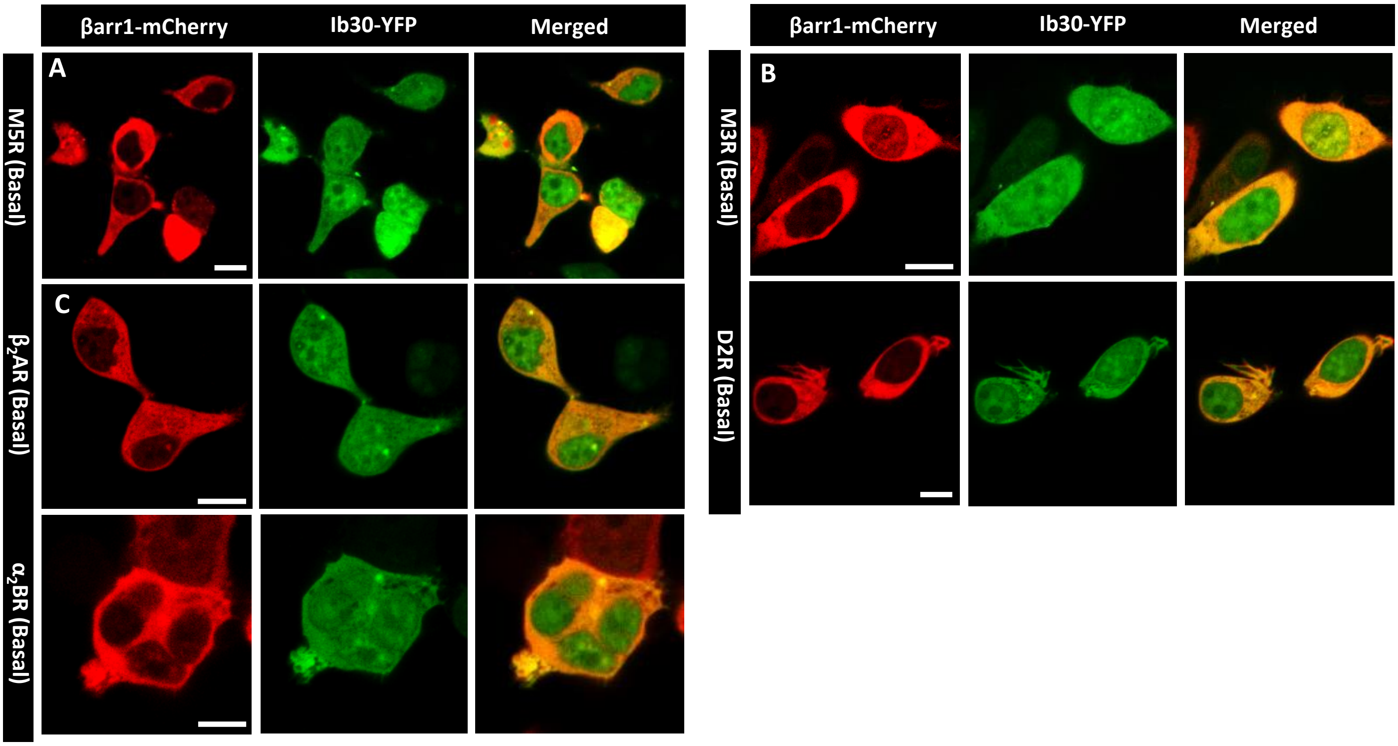


**Supplementary Figure 12. Intrabody30 reports recruitment and trafficking of βarr1 for broad set of native Class A GPCRs.** HEK-293 cells expressing the respective receptor, βarr1-mCherry and Ib30-YFP were stimulated with agonist for indicated time-points and the localization of βarr1 and Ib30 were visualized using confocal microscopy. Scale bar is 10μm.

**Supplementary Figure 13.**

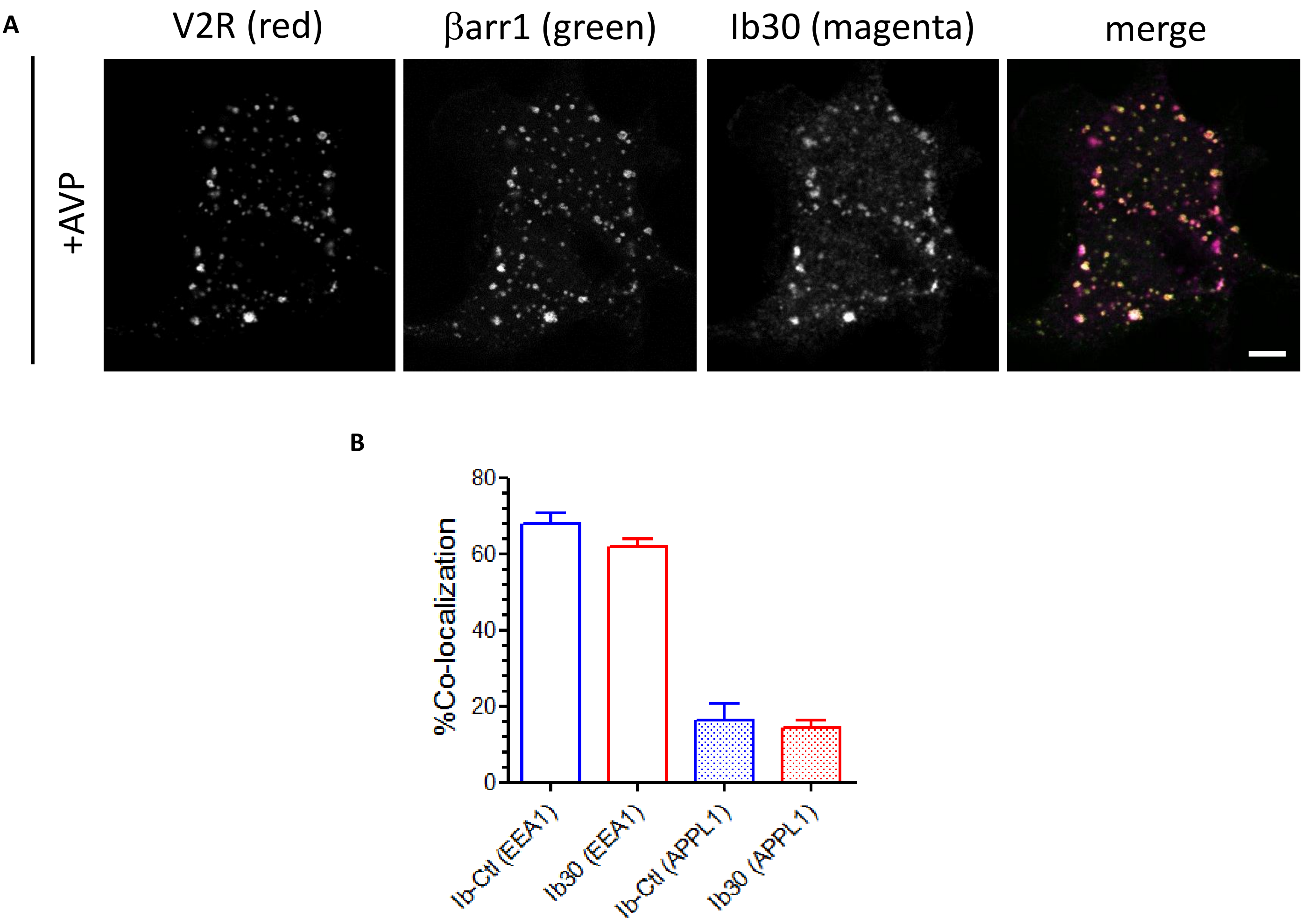


**Supplementary Figure.13. Effect of intrabodies on receptor endocytosis. A** Representative image of Ib30 (magenta), βarr1 (green) and $V_2R$ (red) showing localization of βarr1 mediated endocytosis of $V_2R$ in response to agonist stimulation (AVP 100 nM) as assessed by TIRF microscopy. B. Quantified data represents co-localization of V2R with two early endosomal marker EEA1 and APPL1 with TIRF microscopy.

**Supplementary Figure 14.**

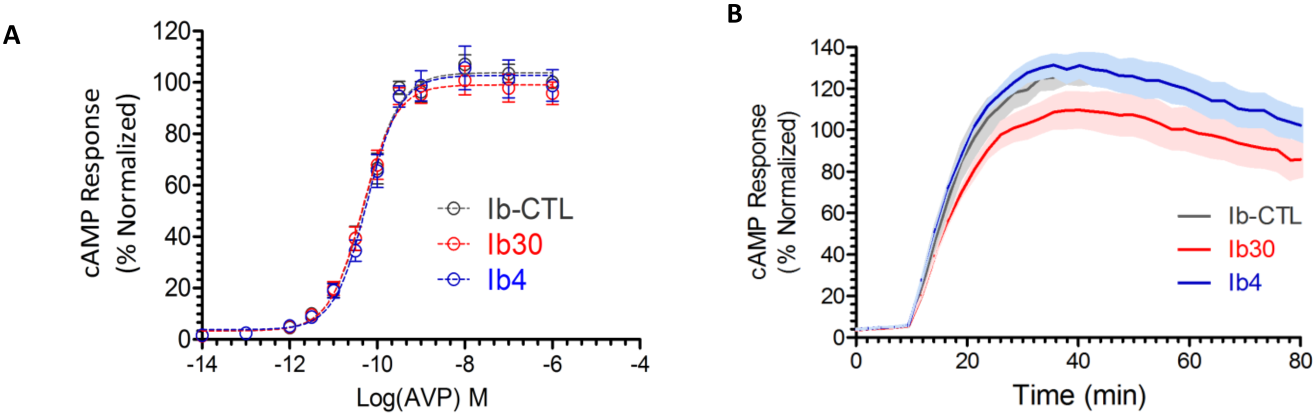


**Supplementary Figure 14. Effect of intrabodies on G protein coupling. A.** HEK-293 cells expressing the $V_2R$, intrabodies (Ib30/Ib4) and a luciferase-based cAMP biosensor were stimulated with varying doses of AVP and levels of cAMP were determined by the bioluminescence generated using a microplate reader. Data are normalized with respect to the maximal response obtained in presence of the control Intrabody (Ib-CTL) and the graph represents average ± s.e.m from three independent experiments. **B.** Graph shows time-kinetics of cAMP response in presence of intrabodies. The experiment was performed in a similar fashion as in panel B except that cAMP levels were recorded in a continuous manner after stimulation of cells with saturating concentration of AVP(100 nM).

**Supplementary Figure 15.**

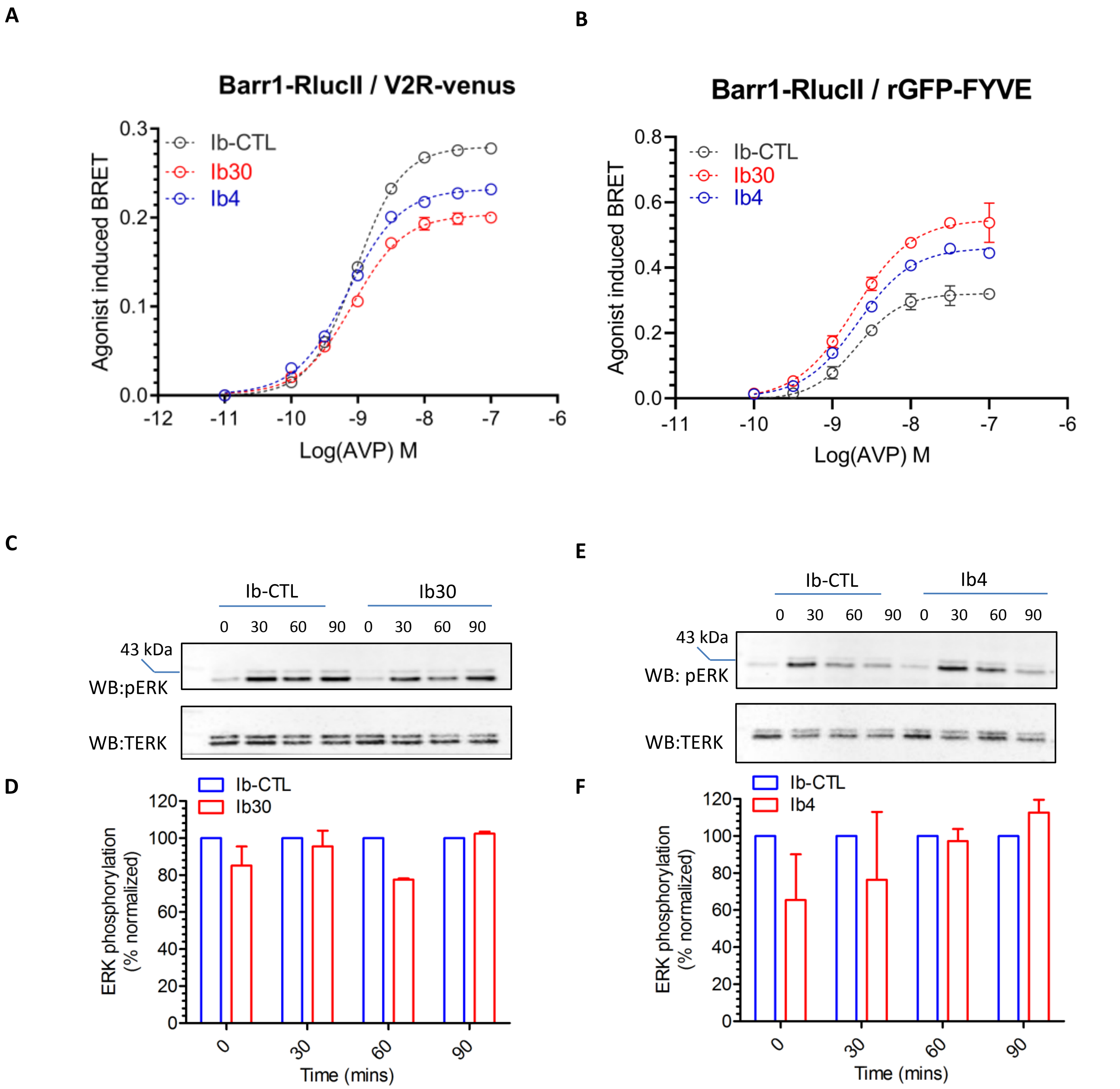


**Supplementary Figure 15. Effect of intrabodies on βarr1 mediated endocytosis & MAPK signaling. A-B** Intrabodies slightly enhance βarr1 recruitment as assessed by intermolecular BRET experiments. HEK-293 cells expressing either the $V_2$R-venus, βarr1-RlucII and intrabodies or the βarr1-RlucII and rGFP-FYVE and intrabodies were stimulated with varying doses of AVP and levels of BRET were recorded using a plate reader. Data represents average ± s.e.m from three independent experiments. **C and E** HEK-293 cells were transfected with FLAG-tagged $V_2$R and Ib30 or Ib4 (or Ib-CTL). 48h post-transfection , cells were stimulated with agonist (AVP 100 nM) at the indicated time points to measure ERK activation through only βarr component of GPCR signaling. ERK phosphorylation was visualized by Immunoblotting. Image shown is representative of two independent experiments. **D and F** Graph shows densitometry based quantification of the data and has been normalized by treating ERK phosphorylation in presence of Ib-CTL as 100% for every indicated time-points. Data represents mean ± s.e.m. of two independent experiments and has been analyzed by using two-way ANOVA with Bonferroni post-test.

**Supplementary Figure 16.**

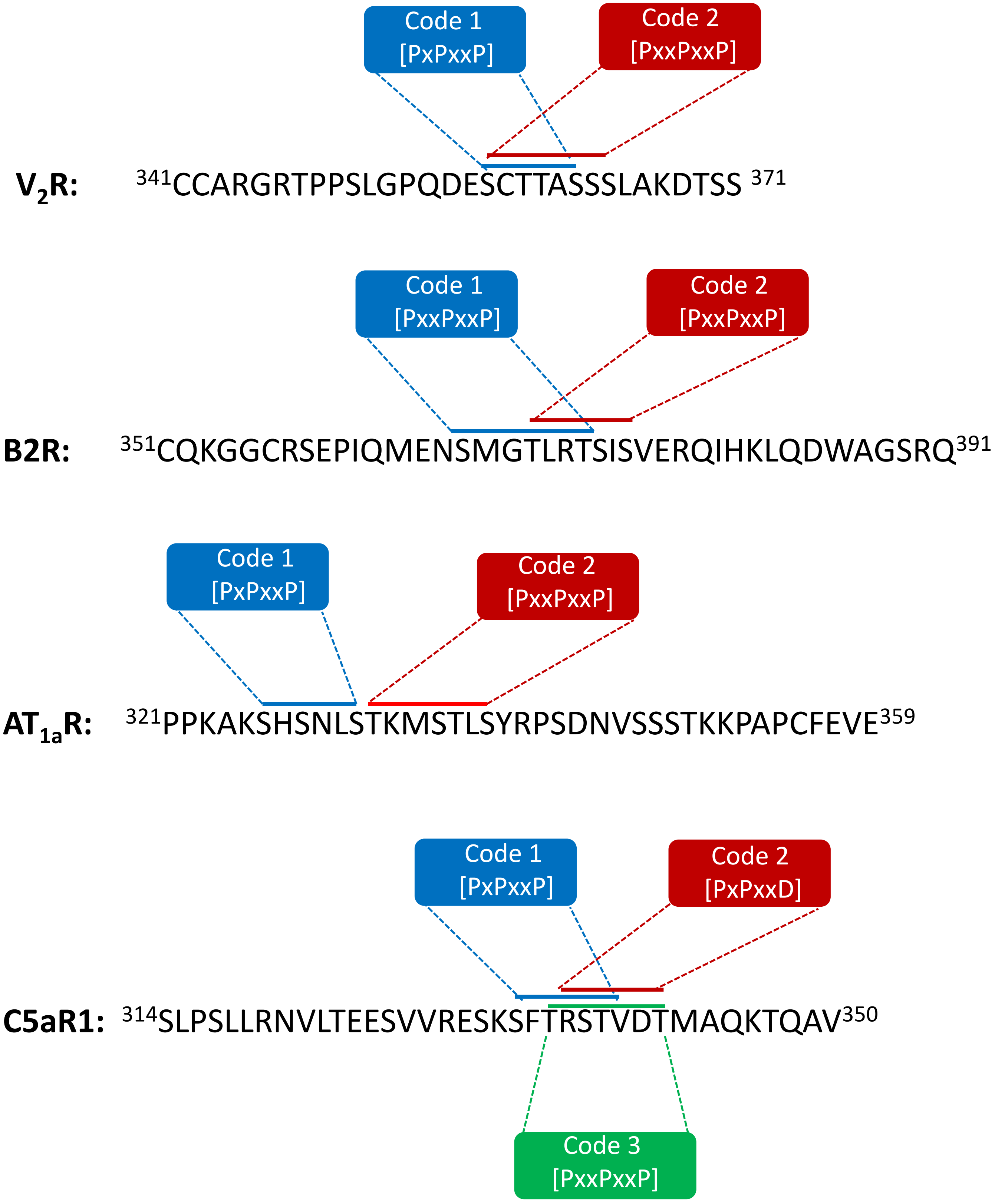


**Supplemental Figure 16. Comparison of full phosphorylation codes in selected GPCRs.** Primary sequence of the carboxyl-terminus of V2R, B2R, $AT_{1a}R$ and C5aR1 is analyzed to identify and highlight full phosphorylation codes based on a previously proposed phospho-code signature by Xu et al., 2017.

**Supplementary Figure 17.**

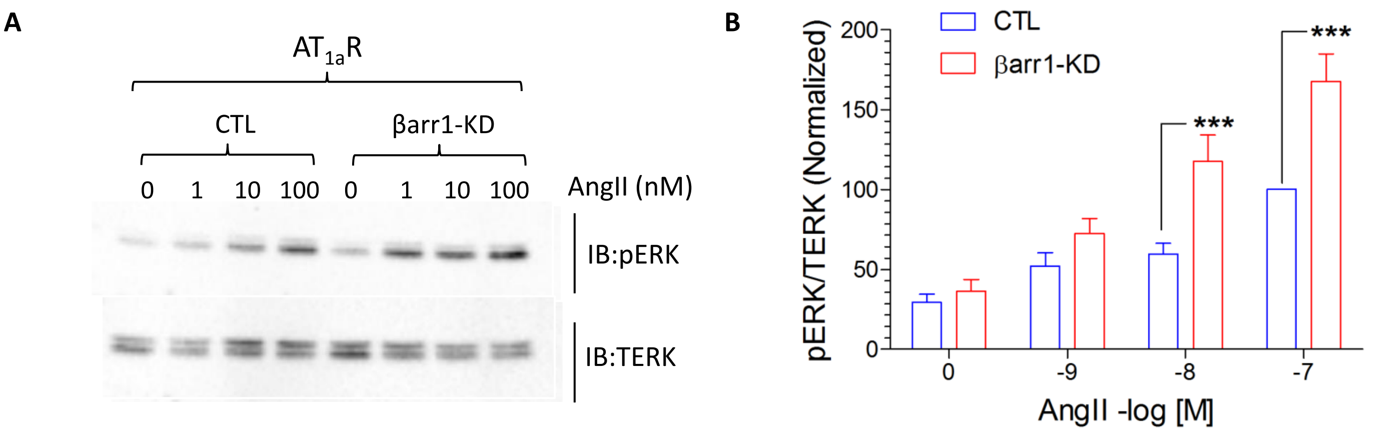


**Supplementary Figure 17. A-F Distinct βarr1 conformations are linked to ERK1/2 MAP kinase phosphorylation.** Agonist-induced phosphorylation of ERK1/2 in HEK-293 cells expressing the indicated receptor in presence and absence of βarr1 knock-down are measured using Western blotting for A. $V_2R$ C. B2R and E. $AT_{1a}R$. Densitometry-based quantification of data is presented as bar-graphs in the right panels. Graph represents mean ± s.e.m. of four independent experiments for $V_2R$, four independent experiments for B2R and five independent experiments for $AT_{1a}R$. For normalization highest concentration of agonist in control cells have been treated as 100% and data has been normalized by two-way ANOVA with Bonferroni post-test (***P<0.001, **P<0.01).

**Supplementary Figure 18.**

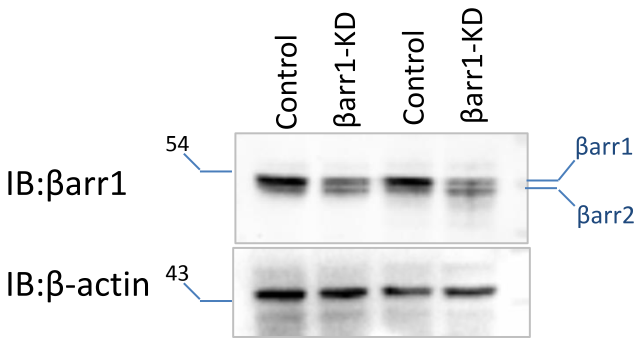


**Supplemental Figure 18. Validation of βarr1 knock-down.** HEK-293 cells stably expressing either control or βarr1-shRNA were lysed, and cellular lysate was subjected to Western blotting. βarrs were detected using anti-βarr antibody and actin levels in cellular lysates were used as loading control.

**Supplementary Figure 19**

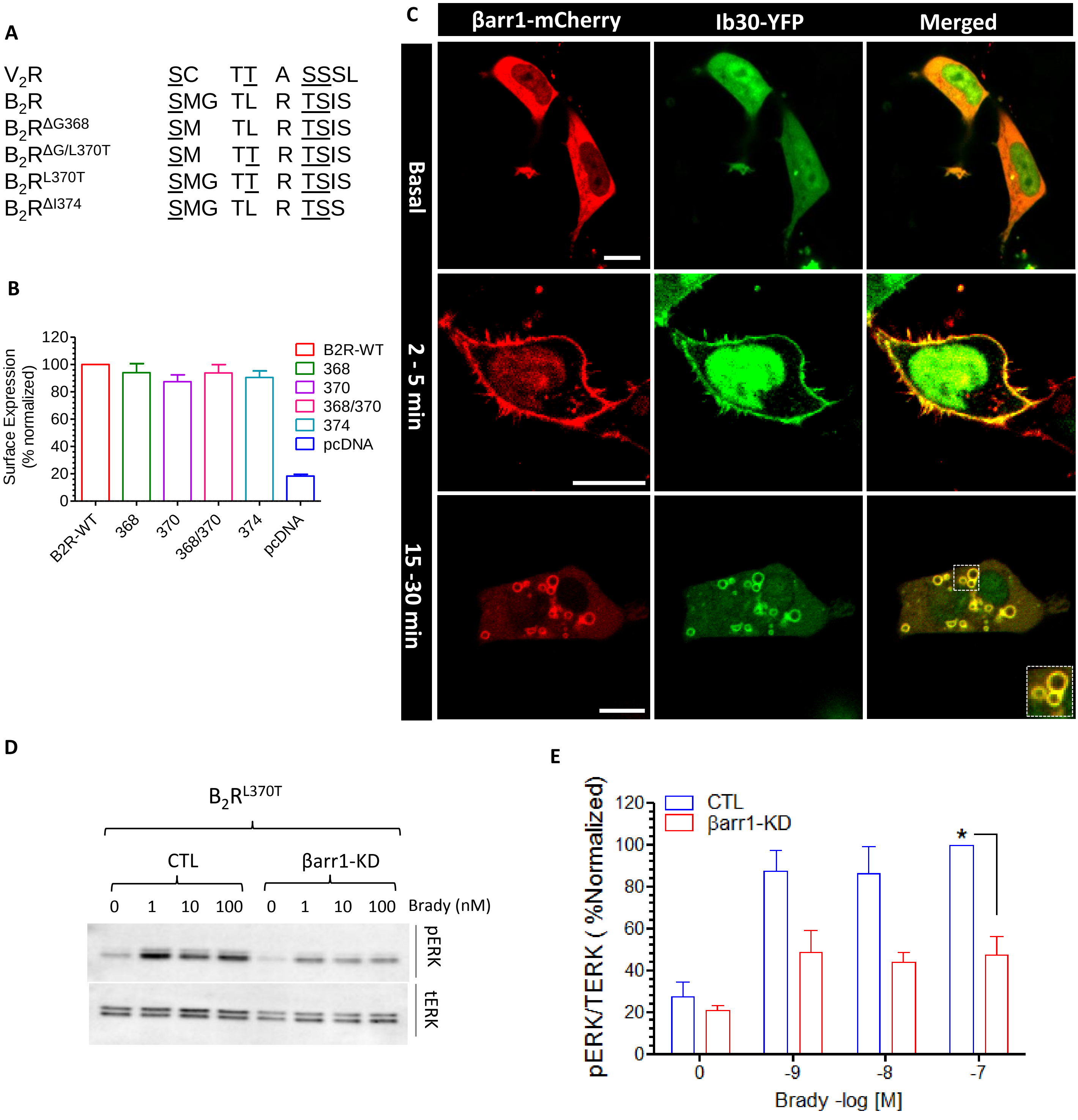


**Supplemental Figure 19. Spatial distribution of phosphorylation sites determine βarr1 conformation and its functional contribution in ERK1/2 phosphorylation. A.** Mutants of B2R generated to decipher the contribution of specific phosphorylation sites determining Fab30 reactivity and ERK1/2 activation. **B.** Quantification of surface expression of B2R and its mutants, as measured by cell based ELISA. **C.** in βarr1 HEK-293 cells expressing $B_2R^{L370T}$, βarr1-mCherry and Ib30-YFP were stimulated with agonist for indicated time-points and the localization of βarr1 and Ib30 were visualized using confocal microscopy. Scale bar is 10µm. **D-E.** Agonist-induced phosphorylation of ERK1/2 in HEK-293 cells expressing $B_2R^{L370T}$ in presence and absence of βarr1 knock-down are measured using Western blotting (left panel) . Densitometry-based quantification of data from three independent experiments is presented as bar-graphs in the right panels normalized with respect to maximal signal under control condition (treated as 100%) and analyzed using One-way-ANOVA.

**Supplementary Figure 20.**

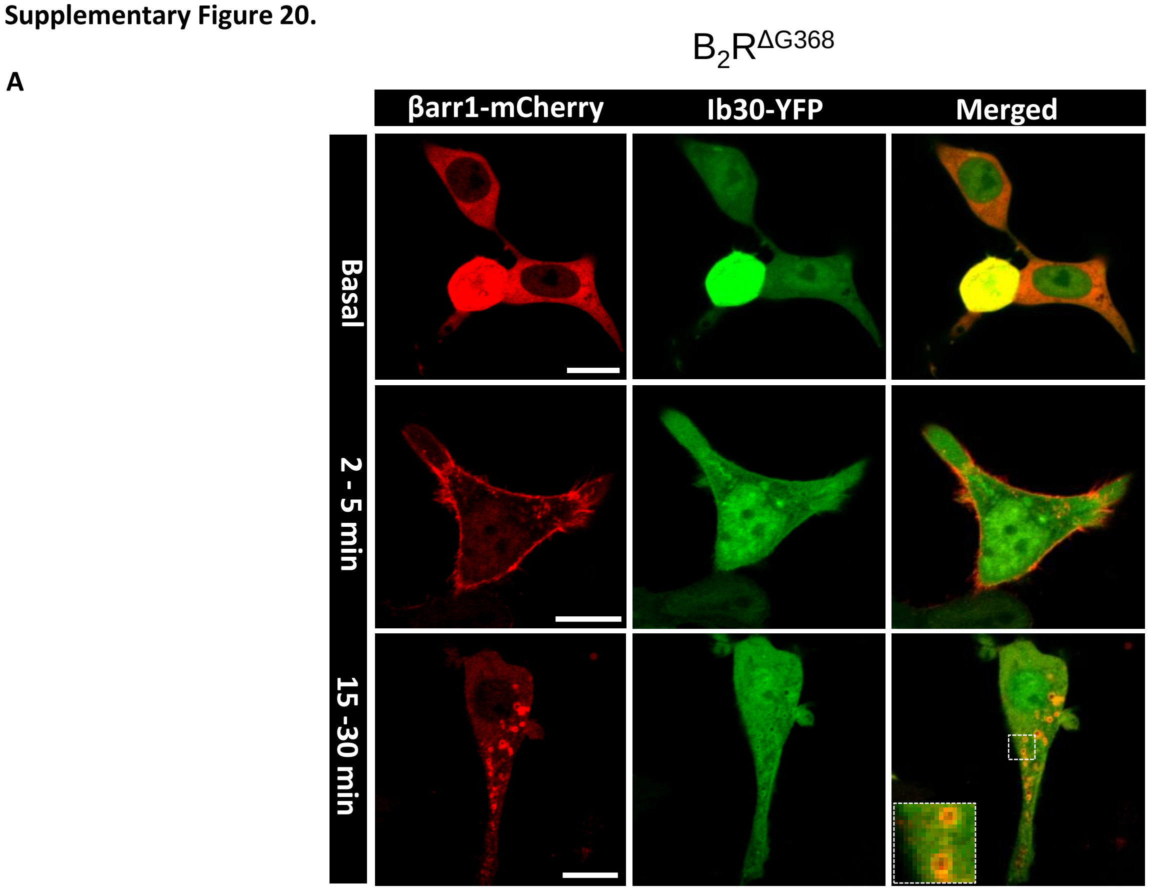


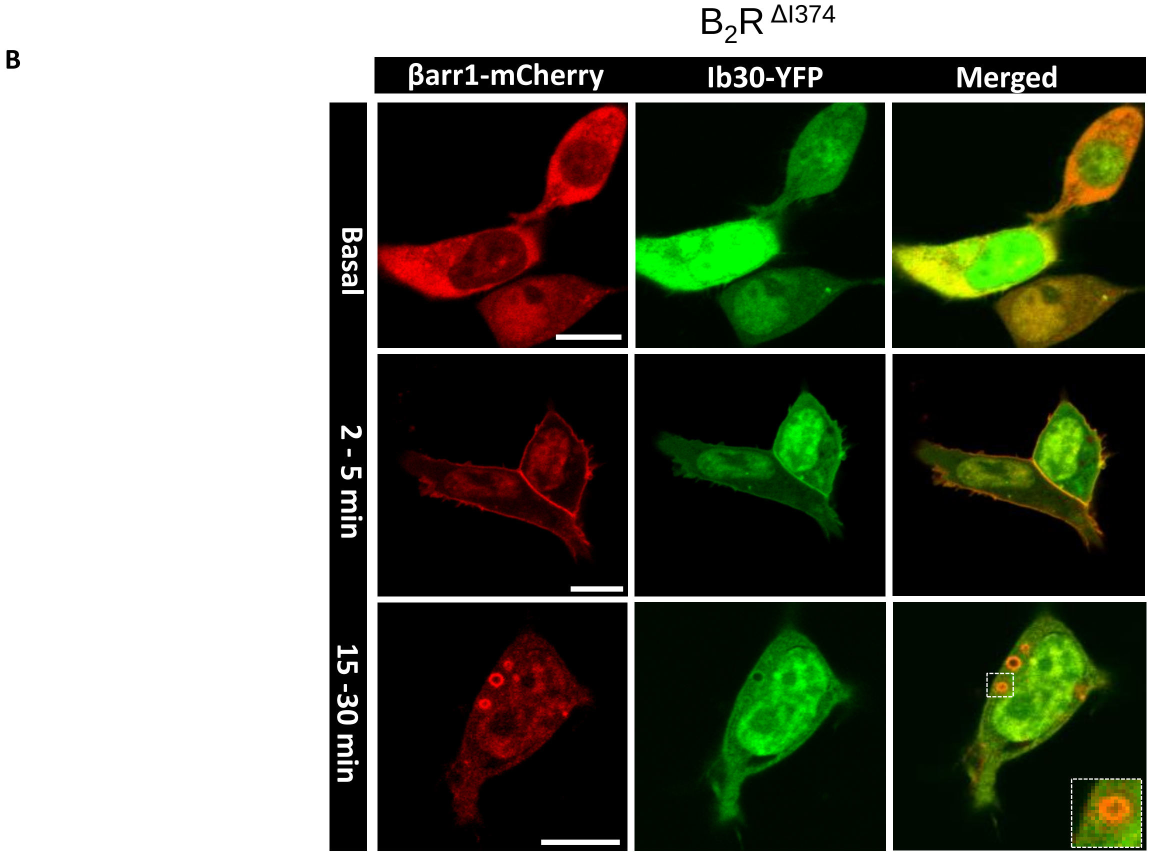


**Supplemental Figure 20. Supplemental Figure 19. Spatial distribution of phosphorylation sites determine βarr1 conformation and its functional contribution in ERK1/2 phosphorylation.** HEK-293 cells expressing $B_2R^{\Delta G368}$ (**A**) or $B_2R^{\Delta I374}$ (**B**) βarr1-mCherry and Ib30-YFP were stimulated with agonist for indicated time-points and the localization of βarr1 and Ib30 were visualized using confocal microscopy. Scale bar is 10μm.

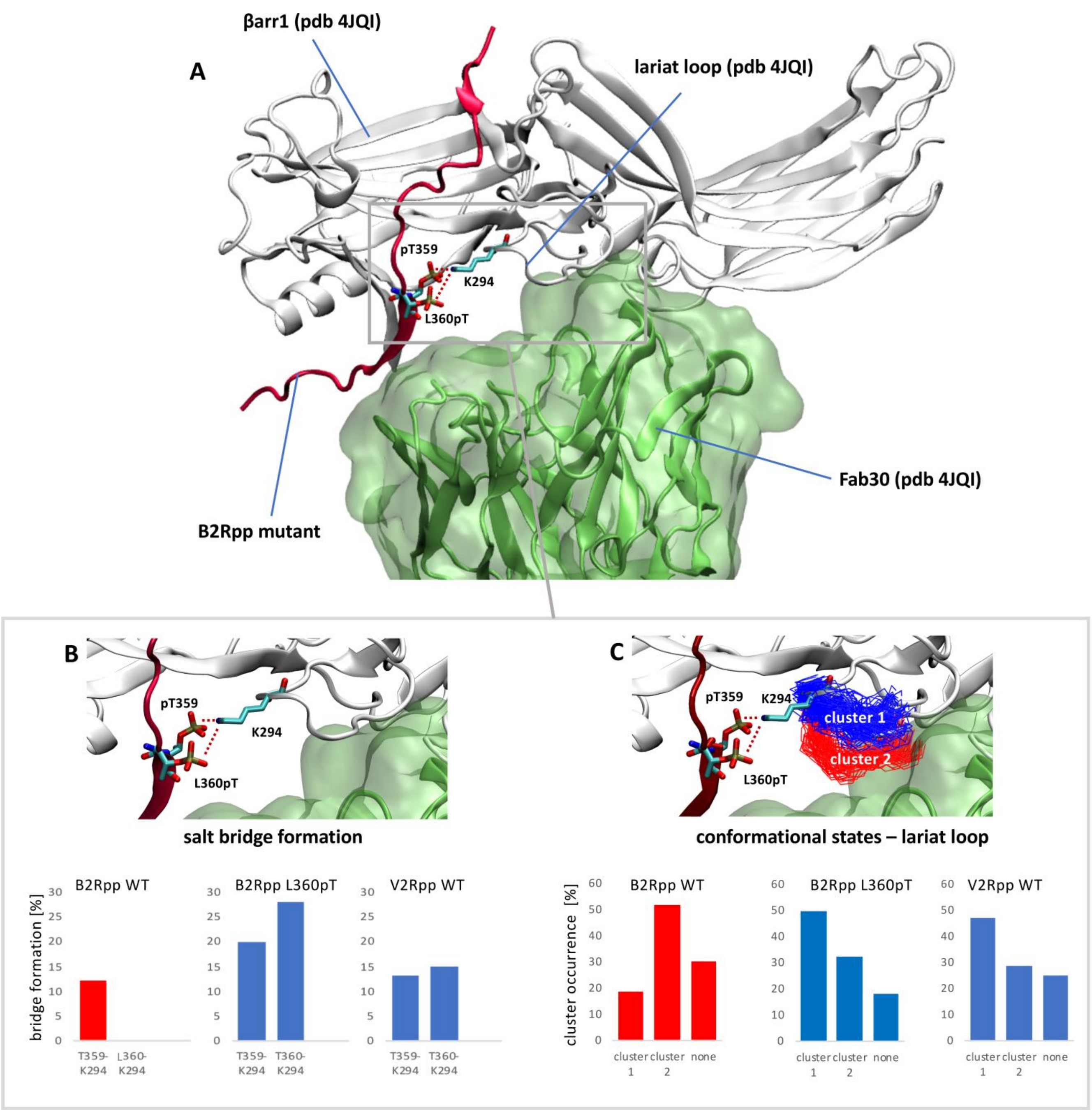


**Supplemental Figure 21.** The phosphorylated receptor C-tail drives Fab30 recognition in the βarr1 complex. (A) Complex of the phosphorylated B2R C-tail containing a L360pT mutation (B2Rpp, red cartoon) with βarr1 (grey cartoon, PDB: 4JQI) and Fab30 (green cartoon and surface, PDB: 4JQI). A bifurcated salt bridge links the B2Rpp (pT359 and L360pT) to the lariat loop (K294) which forms part of the Fab30 binding epitope. (B) Salt bridge formation between phosphorylated receptor C-tail and the lariat loop. The frequency of salt bridge formation was computed over 4µs accumulated simulation time per complex using a distance threshold of 3.2 Å between the oxygens of the phosphate group of phosphorylated threonines and the protonated nitrogen of K294. The L360pT mutant of B2Rpp and the V2Rpp WT establish a bifurcated interaction via pT359 and pT360 with the lariat loop (K294) (blue bar plots). In contrast, B2Rpp WT forms only a single salt bridge as a non-polar residue leucine is present in position 360 (red bar plot). (C) Differential lariat loop stabilization correlates with distinct conformational states. Simulations of 4µs per complex were clustered with rmsd cutoff of 2.2 Å yielding cluster 1 and 2. "None" reflects conformations that did not fulfill this condition. The L360pT mutant of B2Rpp and the V2Rpp WT with a conserved bifurcated salt bridge adopt preferentially conformational states belonging to cluster 1 (blue line depiction in structures and blue bar plot). In contrast, the B2Rpp WT with a single salt bridge favors loop conformations of cluster 2. Cluster 2 is characterized by a downward shift of the lariat loop in the Fab30 recognition interface compared to cluster 1.
